# Rhizospheric and endophytic *Pseudomonas aeruginosa* in edible vegetable plants share molecular and metabolic traits with clinical isolates

**DOI:** 10.1101/2021.06.11.448042

**Authors:** Sakthivel Ambreetha, Ponnusamy Marimuthu, Kalai Mathee, Dananjeyan Balachandar

**Affiliations:** Department of Agricultural Microbiology, Tamil Nadu Agricultural University, Coimbatore, Tamil Nadu, India; Department of Human and Molecular Genetics, Herbert Wertheim College of Medicine, Florida International University, Florida, United States of America; Biomolecular Sciences Institute, Florida International University, Miami, FL, USA

**Keywords:** ERIC PCR, BOX PCR, Auxin, Plant growth-promoting rhizobacteria, PGPR, Agricultural P. *aeruginosa*

## Abstract

*Pseudomonas aeruginosa,* a leading opportunistic pathogen causing hospital-acquired infections is predominantly present in agricultural settings. There are minimal attempts to examine the molecular and functional attributes shared by agricultural and clinical strains of *P. aeruginosa.* This study aims to investigate the presence of *P. aeruginosa* in edible vegetable plants (including salad vegetables) and analyze the evolutionary and metabolic relatedness of the agricultural and clinical strains. Eighteen rhizospheric and endophytic *P. aeruginosa* strains were isolated from cucumber, tomato, eggplant, and chili directly from the farms. The identity of these strains was confirmed using biochemical, and molecular markers and their genetic and metabolic traits were compared with clinical isolates. DNA fingerprinting analyses and 16S rDNA-based phylogenetic tree revealed that the plant- and human-associated strains are evolutionarily related. Both agricultural and clinical isolates possessed plant-beneficial properties, including mineral solubilization (phosphorous, potassium, and zinc), ammonification, and the ability to release extracellular siderophore and indole-3 acetic acid. These findings suggest that rhizospheric and endophytic *P. aeruginosa* strains are genetically and functionally analogous to the clinical isolates. This study highlights the edible plants as a potential source for human and animal transmission of *P. aeruginosa*.

## Introduction

*Pseudomonas aeruginosa* was discovered in 1882 when Carle Gessard noticed a bluish green color in injured soldiers’ bandages (Gessard 1984). This color was due to the blue-green phenazine compound synthesized by *P. aeruginosa,* making it distinct from other Pseudomonads (Reyes et al. 1981; Turner and Messenger 1986). *P. aeruginosa* is a Gram-negative gamma-proteobacterium that colonizes diverse host systems, including the nematodes, insects, plants, animals, and humans (Schroth et al. 1977; Botzenhart and Doring 1993; Banerjee and Dangar 1995; Rahme et al. 1995). Due to its recalcitrant pathogenicity and drug resistance, this omnipresent organism is listed as a serious threat pathogen by the US Centers for Disease Control (CDC), World Health Organization (WHO), and UK Public Health England (PHE) (CDC AR, 2019; WHO News, 2019; PHE, 2020)

*P. aeruginosa* causes fatal infections in immunocompromised individuals and patients with genetic disorders such as cystic fibrosis (Reynolds et al. 1975; Von Graevenitz 1977). Its infections in otherwise healthy individuals include folliculitis, endocarditis, osteomyelitis, and sclerokeratitis (Radford et al. 2000; Tate et al. 2003; Doustdar et al. 2019). According to the International Nosocomial Infection Control Consortium (INICC), *P. aeruginosa* is among the significant hospital-acquired pathogens leading to ventilator-, surgical implant-, central line- and urinary catheter-associated infections (Rosenthal et al. 2020). *P. aeruginosa* releases an arsenal of virulence factors that include rhamnolipid, pyocyanin, pyoverdine, pyochelin, elastases, proteases, lipases, polysaccharides, hydrogen cyanide, and exotoxins to breach the mucus barriers and establish the infection (Balasubramanian et al. 2012; Moradali et al. 2017).

The role of *P. aeruginosa* in the agricultural ecosystem is controversial. Several studies showed that *P. aeruginosa* supports plant growth and disease control through multiple mechanisms (Ali Siddiqui and Ehteshamul-Haque 2001; Adesemoye and Ugoji 2009; Yasmin et al. 2014; Radhapriya et al. 2015; Arif et al. 2016; Durairaj et al. 2017; Gupta and Buch 2019; Chandra et al. 2020). *P. aeruginosa* helps in the solubilization of complex soil minerals (tri-calcium phosphate, potassium aluminum silicate, and zinc oxide), ammonification, and nitrification (Obaton et al. 1968; Illmer and Schinner 1992; Fasim et al. 2002; Jha et al. 2009; Gupta and Buch 2019). This bacterium also releases salicylic acid, hydrogen cyanide, pyocyanin, rhamnolipid, and siderophores to inhibit the growth of other pathogens and insect pests that compete with it in the agricultural ecosystem (Kloepper et al. 1980; Cartwright et al. 1995; De Meyer and Höfte 1997; Kim et al. 2000; Audenaert et al. 2002). Conversely, some argue that *P. aeruginosa* is a plant pathogen that inhibits seed germination and causes rot and wilt in maize, ginseng, melon, chickpea, and tobacco (Clara 1930; Elrod and Braun 1942; Mondal et al. 2012; Gao et al. 2014; Tiwari and Singh 2017). It was believed that *P. aeruginosa* colonizing humans and plants were two different species in the early days. Schroth and his associates proved that the clinical strains of *P. aeruginosa* could also cause plant infections (Schroth et al. 1977). His group also identified that the most used virulent strain *P. aeruginosa* PA14, isolated initially from Pittsburgh burn ward, causes extensive plant rot in cucumber, lettuce, potato, and tomato (Mathee 2018; Schroth et al. 2018).

To date, *P. aeruginosa* has been reported in tomato, radish, celery, carrot, endive, cabbage, onion, lettuce, watercress, chicory, Swiss chard, and cucumber from the hospital kitchens in the USA and Brazil (Kominos et al. 1972; Wright et al. 1976; Correa et al. 1991). Furthermore, the *P. aeruginosa* contamination in fresh vegetables have been reported in the retail markets, supermarkets, local vendors, and canteens in India, Jamaica, France, Germany, Ireland, Holand, and United Kingdoms (Viswanathan and Kaur 2001; Curran et al. 2005; Allydice-Francis and Brown 2012). Agricultural soil and plants as the source of *P. aeruginosa* infection was first reported in 1970’s (Green et al. 1974; Cho et al. 1975). However, Deredjian et al. (2014) reported a low occurrence of *P. aeruginosa* in agricultural soil in France and Burkina Faso. Most of the studies on agricultural *P. aeruginosa* have focused on characterizing one or two strains (Kumar et al. 2013; Jasim et al. 2014; Akinsanya et al. 2015; Shi et al. 2015; Devi et al. 2017; Wu et al. 2018; Roychowdhury et al. 2019; Iasur Kruh et al. 2020; Mukherjee et al. 2020; Singh et al. 2021; Sun et al. 2021). There is a clear gap in looking into the vegetable plants’ rhizosphere and their internal tissues as the potential source of *P. aeruginosa*. In the current study, we have isolated *P. aeruginosa* from the rhizosphere and internal tissues of the vegetable plants directly from the farms in Southern India. Their genetic and metabolic characteristics were compared with well-characterized clinical *P. aeruginosa*.

## Materials and Methods

### Bacterial strains and culture conditions

Clinical strains of *P. aeruginosa*, PAO1, ATCC10145, and ATCC9027 were used as controls (Haynes 1951; Holloway 1955; Picard et al. 1994). A well-characterized plant growth-promoting rhizobacteria (PGPR), *Bacillus altitutinis*, FD48 (Table 1) was used as the control for all PGPR experiments. Commercial phosphorous solubilizing bacteria *Bacillus megaterium* strain var *phosphaticum* Pb1, potassium releasing bacteria *Bacillus mucilaginous* strain KRB9, and zinc solubilizing bacteria *Pseudomonas chloraraphis* strain ZSB15 (Table 3) were used as the positive controls for estimating the respective mineral solubilization experiments. *P. aeruginosa* strains were periodically sub-cultured and grown in Pseudomonas agar (for pyocyanin) medium (PAP, Himedia); *P. chloraraphis* in King’s B medium (King et al. 1954); *B. altitudinis* and *B. mucilaginous* in nutrient agar medium at 37°C.

**Table 1.** Number of *P. aeruginosa* isolates screened in this study.

| **Ecosystem** | **Crop** | **Initial No. of isolates** | **No. of isolates**<br>**Morphological screening** | **No. of isolates**<br>**Molecular screening** |
| --- | --- | --- | --- | --- |
| Orchard | Cucumber rhizosphere | 21 | 8 | 3 |
|  | Cucumber endophyte | 15 | 4 | 1 |
|  | Tomato rhizosphere | 10 | 3 | 2 |
|  | Tomato endophyte | 23 | 6 | 4 |
|  | Eggplant rhizosphere | 35 | 9 | 3 |
|  | Eggplant endophyte | 16 | 7 | 1 |
|  | Chili rhizosphere | 28 | 6 | 1 |
|  | Chili endophyte | 22 | 4 | 3 |
|  | Bottle gourd rhizosphere | 26 | 4 | 0 |
|  | Bottle gourd endophyte | 32 | 2 | 0 |
|  | Bitter gourd rhizosphere | 11 | 1 | 0 |
|  | Bitter gourd endophyte | 15 | 4 | 0 |
| Wetland | Rice rhizosphere | 26 | 2 | 0 |
|  | Rice endophyte | 20 | 3 | 0 |
|  | Total isolates | 300 | 63 | 18 |

### Isolation of plant-associated and rhizosphere *P. aeruginosa*

Samples were collected from edible crops, namely rice, tomato, cucumber, eggplant, chili, and bottle, and bitter gourds and their associated rhizosphere (soil adhering to the roots). These samples were from the wetland and garden land ecosystems of Tamil Nadu Agricultural University, India (latitude, 11° 07’ 3.36“; longitude 76° 59’ 39.91”). In each crop field, five plant samples were collected from random sites and pooled together.

For isolation of rhizospheric *P. aeruginosa*, 10 g of the soil sample was pooled from five crops of each kind. A 100-ml sterile distilled water (pH 7.0) was added to the soil and mixed vigorously. The soil solution was then serially diluted up to 10^-4^. One ml of the diluent combined with 20 ml of PAP medium was plated using the conventional pour plate technique (Van Soestbergen and Lee 1969; Elbadry et al. 1999). The plates were incubated overnight at 37°C.

For isolation of the endophytes, the plant samples were surface sterilized using sodium hypochlorite (5% chlorine) for 10 min to remove epiphytic and saprophytic organisms (Gardner et al. 1982). The samples were then washed with sterile distilled water. The wash water was collected and plated to confirm the absence of any surface microbes. The surface-sterilized plant samples were crushed in sterile pestle and mortar. The crushed samples were mixed with sterile distilled water (pH 7.0) in 1:10 weight/volume. This mixture was serially diluted with sterile distilled water up to 10^-2^. One ml of the diluent combined with 20 ml of PAP medium was plated using the pour plate technique (Van Soestbergen and Lee 1969; Elbadry et al. 1999). The plates were incubated overnight at 37°C.

### Pyocyanin production

The colonies were initially screened based on pyocyanin production, indicative of *P. aeruginosa*, by the bluish-green discoloration on the PAP medium. For further confirmation, selected individual isolates were grown in 30 ml of glycine-alanine broth (Ingledew and Campbell 1969; Devnath et al. 2017). The 48-h cultures were centrifuged at 5000 g for 15 minutes. The supernatant was mixed with 0.5 V chloroform, vortexed, and allowed to settle for 10 min. The blue solvent layer (bottom) was acidified with 0.2 V of 0.1 N HCl, and its absorbance was measured at 520 nm (Varian Cary® −50, Australia) (Essar et al. 1990). The concentration of pyocyanin (µg/ml) was estimated by multiplying of OD_520_ with pyocyanin extinction coefficient (17.072) (Kurachi 1958).

### Molecular screening

Genomic DNA was extracted from the select isolates using the hexadecyl trimethyl ammonium bromide method (Melody 1997). The purity of DNA was analyzed using a Nanodrop spectrophotometer (Thermo Scientific, Nanodrop^TM^ 2000c). The absorbance of 1.8 at 260/280 was used as the indicator of DNA purity. *P. aeruginosa* genus-(PA-GS-F/R) and species- (PA-SS-F/R) specific primers (Table 3) were used for molecular confirmation of the chosen isolates as previously described (Spilker et al. 2004).

### 16S rDNA sequencing

Nearly full-length *P. aeruginosa* 16S rRNA genes were amplified using universal eubacterial primers (Table 2; Weisburg et al. 1991). PCR amplification was performed in a thermocycler (Bio-Rad T-100^TM^, USA). The amplified genes were sequenced in both directions by Sanger’s chain termination method Sanger et al. (1977) using an Applied Biosystems automated sequencer (Bioserve, Hyderabad, India). The 16S rRNA gene sequences of the plant-associated *P. aeruginosa* strains (numbered PPA1 to PPA18) isolated and identified in the current study were submitted to NCBI GenBank (Accession no. MT734694 to MT734711).

**Table 2.**
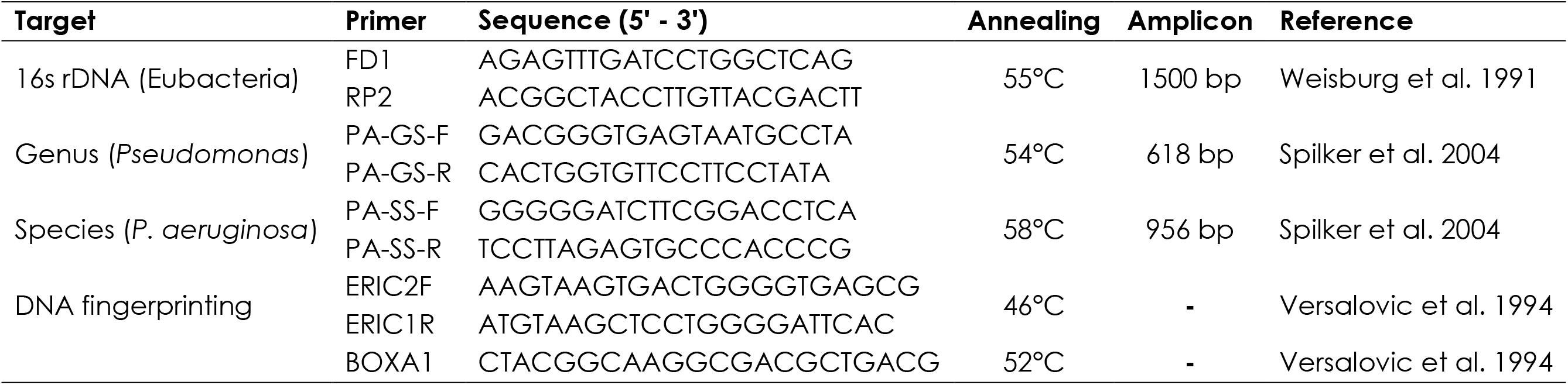
Details of primers used in the study.

### Repetitive element PCR based DNA fingerprinting analysis

Primers complementary to the interspersed repetitive sequences (or repeated DNA elements) conserved within the prokaryotic genome are used for rep-PCR-based fingerprinting (Versalovic et al. 1994). The conserved repetitive elements used for bacterial fingerprinting include Enterobacterial repetitive intergenic consensus (ERIC) and BOX element (BOX). The rep-PCR technique distinguishes the closely related bacterial strains.

#### ERIC fingerprinting

ERIC primers constitute oligonucleotides complementary to the conserved (126-bp) palindromic regions (Table 3; Versalovic et al. 1994). The amplified genomic DNA of the *P. aeruginosa* isolates were separated in 2% agarose gel and visualized their banding patterns in the Gel Doc^TM^ XR^+^ documentation system (Bio-Rad, USA).

**Table 3.** Bacterial strains used in the study.

| **Bacteria** | **Source** | **Infection/Niche** | **References** |
| --- | --- | --- | --- |
| ***Pseudomonas aeruginosa* (reference strains)** |  |  |  |
| PAO1 | Human | Wound infection | Holloway, (1955) |
| ATCC9027 | Human | Otitis externa | Haynes, (1951) |
| ATCC10145 | Human | Unknown | Picard et al. (1994) |
| **Plant beneficial bacteria (reference strains)** |  |  |  |
| *Bacillus altitudinus*, FD48 | Rice | Phyllosphere | Kumar et al. (2017) |
| *Bacillus megaterium* var *phosphaticum*, Pb1 | Soil | Rhizosphere | Balamurugan and Gunasekaran, (1996) |
| *Paenibacillus mucilaginosus*, KRB9 | Soil | Rhizosphere | Brindavathy and Gopalaswamy, (2014) |
| *Pseudomonas chlororaphis* strain, ZSB15 | Rice | Rhizoplane | Bowya and Balachandar, (2020) |
| **Plant-associated *P. aeruginosa* strains** |  |  |  |
| PPA01 | Cucumber | Rhizosphere | This study |
| PPA02 | Cucumber | Rhizosphere | This study |
| PPA03 | Cucumber | Endophyte | This study |
| PPA04 | Cucumber | Rhizosphere | This study |
| PPA05 | Tomato | Endophyte | This study |
| PPA06 | Tomato | Rhizosphere | This study |
| PPA07 | Tomato | Endophyte | This study |
| PPA08 | Tomato | Endophyte | This study |
| PPA09 | Tomato | Rhizosphere | This study |
| PPA10 | Tomato | Endophyte | This study |
| PPA11 | Eggplant | Endophyte | This study |
| PPA12 | Eggplant | Rhizosphere | This study |
| PPA13 | Eggplant | Rhizosphere | This study |
| PPA14 | Eggplant | Rhizosphere | This study |
| PPA15 | Chili | Rhizosphere | This study |
| PPA16 | Chili | Endophyte | This study |
| PPA17 | Chili | Endophyte | This study |
| PPA18 | Chili | Endophyte | This study |

#### BOX fingerprinting

An oligonucleotide primer complementary to the conserved *boxA* (59 bp) segment was used for BOX fingerprinting analyses (Versalovic et al. 1994). The amplified genomic DNA from the PPA strains was resolved using 2% agarose gel and visualized in the Gel Doc^TM^ XR^+^ documentation system (Bio-Rad, USA).

### Mineral solubilization

The PPA strains were tested for their ability to solubilize the complex soil minerals to release the nutrients such as phosphorous, potassium, and zinc (Bunt and Rovira 1955; Sperber 1958; Hu et al. 2006). The cultures were grown in 10 ml LB broth overnight at 37°C. The cell pellets were harvested and washed with 0.2 M phosphate buffer. The OD_660_ was adjusted to 0.5 with sterile water. A 10 µl of the culture was spot inoculated into selective media with insoluble mineral sources.

Tri-calcium phosphate (Sperberg’s apatite medium), potassium aluminum silicate (Alexandrov’s medium), and zinc oxide (Bunt and Rovira medium) were used to test the phosphorous, potassium, and zinc solubilization potential, respectively (Bunt and Rovira 1955; Sperber 1958; Hu et al. 2006). The plates were incubated for 48 h at 37°C and measured the diameter of the bacterial colonies and their inhibitory (halo) zones. The ability of our strains to solubilize the complex minerals was presented as solubilization index: Ratio of total diameter (colony + halo zone) to colony diameter (Edi-Premono et al. 1996).

### Ammonification

The ability of the isolates to fix atmospheric N into ammonia was tested using Nessler’s reagent (Cappuccino and Sherman 1983; Goswami et al. 2014). Nessler’s reagent consists of potassium tetraiodomercurate and potassium hydroxide which form an insoluble brown precipitate in the presence of ammonia.

The strains were inoculated in 2 ml of peptone broth and incubated at 37°C for 5 days under 100 rpm shaking. The culture supernatant was mixed with an equal volume of Nessler’s reagent and vortexed. Absorbance (A_450_) of the brown insoluble mixture formed was measured using an ELISA reader (Spectramax® i3x, USA). The same procedure was repeated to generate the standard curve with ammonium sulfate (2 mM to 10 mM). The concentration of ammonia (in mM) produced by individual strains was estimated by comparing against the standard curve.

### Indole acetic acid

The amount of indole-3 acetic acid (IAA) released into the medium was detected using Salkowski’s reagent, which is a mixture of 35% perchloric acid and 5M ferric chloride (Gordon and Weber 1951). IAA reduces ferric compound resulting in pink coloration.

The bacterial strains were inoculated into 10 ml LB broth supplemented with 0.2 % L-tryptophan and incubated at 37°C under 125 rpm shaking for 7 days. The culture supernatant was mixed with two volumes of Salkowski’s reagent, and the absorbance (A_530_) was quantified in a multi-mode microplate reader (Spectramax® i3x, USA). The same procedure was repeated to generate the standard curve with IAA (5 ppm to 100 ppm). The concentration of IAA produced by individual strains was determined by comparing against the standard curve.

### Siderophore

#### Qualitative assay

Siderophore production was qualitatively detected using chrome azurol S (CAS) agar medium (Schwyn and Neilands 1987). The CAS agar medium is made of chrome azurol S, hexadecyltrimethylammonium bromide, and iron(III) (colored dye-iron complex). The bacterial siderophores remove the iron(III) from this complex (iron chelation), leading to yellow coloration. All the strains were streaked on CAS agar medium and incubated overnight at 37°C. The formation of yellowish-orange zones around the colony is indicative of siderophore production.

#### Quantitative assay

The CAS-shuttle assay was performed for the siderophore estimation (Schwyn and Neilands 1987). The strains were grown overnight in succinate broth at 37°C. The cell-free supernatant was mixed with an equal volume of CAS solution and incubated at room temperature for 1 h. The absorbance (A_630_) was measured using a spectrophotometer (Varian Cary® - 50, Australia). The percentage of siderophore is based on the equation [(A_r_ -A_s_)/A_r_] x 100, where A_r_ is the A_630_ of reference (CAS assay solution and uninoculated media) and A_s_ is the A_630_ of the sample (CAS assay solution and culture supernatant) (Pérez-Miranda et al. 2007).

### Statistical analysis

All data were subjected to a one-way analysis of variance (ANOVA) with a P-value of 0.05 and Duncan’s multiple range test was performed between individual means to reveal the significant difference (XLSTAT, version 2010.5.05 add-in with Windows Excel). Principal coordinate analysis (PCoA) based on Euclidean distance was carried out in NCSS 2020 statistical software (NCSS, Kaysville, USA). Primer 7 (Plymouth Routines in Multivariate Ecological Research, version 7; PRIMER-E, Plymouth, UK) was employed for non-metric multidimensional scaling of DNA fingerprints based on the Bray-Curtis similarity matrix (Clarke 1993). Data analysis and scientific graphing were done in OriginPro version 8.5 (OriginLab®, USA).

### Bioinformatics analysis

The identity of 16S rRNA gene sequences of the PPA strains isolated in this study was determined by performing a BLASTN similarity search against PAO1 in the Pseudomonas database (https://pseudomonas.com/blast/setnblast; Winsor et al. (2016). The 16S rDNA sequence of ten plant-associated *P. aeruginosa* strains, KSG (Accession no: LN874213), MML2212 (Accession no: EU344794), KKRB-P1 (Accession no: MW149279), MP1 (Accession no: MT937234), Ld-08 16S (Accession no: MT472133), VL4 (Accession no: MN611376), AT5 (Accession no: MN636767), PA4 (Accession no: MN636761), SEGB6 (Accession no: MN565979), and choltrans (Accession no: MK782058) and three clinical strains, PAO1 (Accession no: MT337602), ATCC10145 (Accession no: NR_114471), and ATCC9027 (Accession no: NZ_PDLX0000000) were retrieved from NCBI Genbank (https://www.ncbi.nlm.nih.gov/genbank/). Molecular Evolutionary Genetics Analysis (MEGA) 7.0 software was used for the constructing a phylogenetic tree (Kumar et al., 2016). The 16S rDNA sequences were used to create the tree and their evolutionary history was inferred using the Neighbor-Joining method (Saitou and Nei 1987). Their evolutionary distances were computed using the Maximum Composite Likelihood method (Tamura et al. 2004).

## Results

### Isolation and screening

Human transmission of *P. aeruginosa* through the consumption of raw vegetable salads was first reported in the ‘70s from the USA, followed by a single report in the ‘90s from Brazil (Kominos et al. 1972; Wright et al. 1976; Correa et al. 1991). The current research was undertaken to fill this three-decade gap in testing the edible crops as the source of *P. aeruginosa*. The crops tested in this study include rice, tomato, chili, and cucumber (salad vegetables), eggplant, and gourds. Samples from the rhizosphere and endophyte were plated in triplicates (for a total of 48 plates, 300 CFU/plate) in *P. aeruginosa* selective medium (Table 1). Of these, 300 putative *P. aeruginosa* isolates were sub-cultured in the same medium (data not shown). Out of 300, 63 isolates were selected based on the development of bluish green pigmentation indicative of pyocyanin production, a *P. aeruginosa* biomarker (Gessard 1984; Alatraktchi et al. 2020).

### Genus- and species-level identification

The selected isolates were further screened using *P. aeruginosa* genus- (PA-GS-F/R) and species- (PA-SS-F/R) specific primers that would result in amplicons of 618 bp and 956 bp, respectively (Table 2; Spilker et al. 2004). As expected, the three well-characterized *P. aeruginosa* strains, PAO1, ATCC10145, and ATCC9027, amplified 618 bp (Lane 19-21, respectively; Fig. 1A) and 956 bp (Lane 19-21, respectively; Fig. 1B) fragments with genus- and species-specific primers, respectively. As expected, there was no amplification in the controls, with no template (Lane 23; Fig 1A and B) and unrelated species *Enterobacter cloacae* (Lane 22; Fig 1A and B). After repeating the experiments for three times, out of the 63 isolates selected based on pyocyanin production (as observed on the plates), only 18 strains (29%) were amplifiable using *P. aeruginosa* genus- and species-specific primers (Lanes 1 to18, Fig. 1A and B). The confirmed strains were from the rhizosphere and inner tissues (endophytes) of four plants (tomato, chili, cucumber, and eggplant) (Table 1). Henceforth these strains are referred to as plant-associated *P. aeruginosa* (PPA) 01 to 18 (from cucumber (PPA01 to 04), tomato (PPA05 to 10), eggplant PPA11 to PP14, and chili (PPA15 to PP18)) (Table 3).

**Fig. 1.**
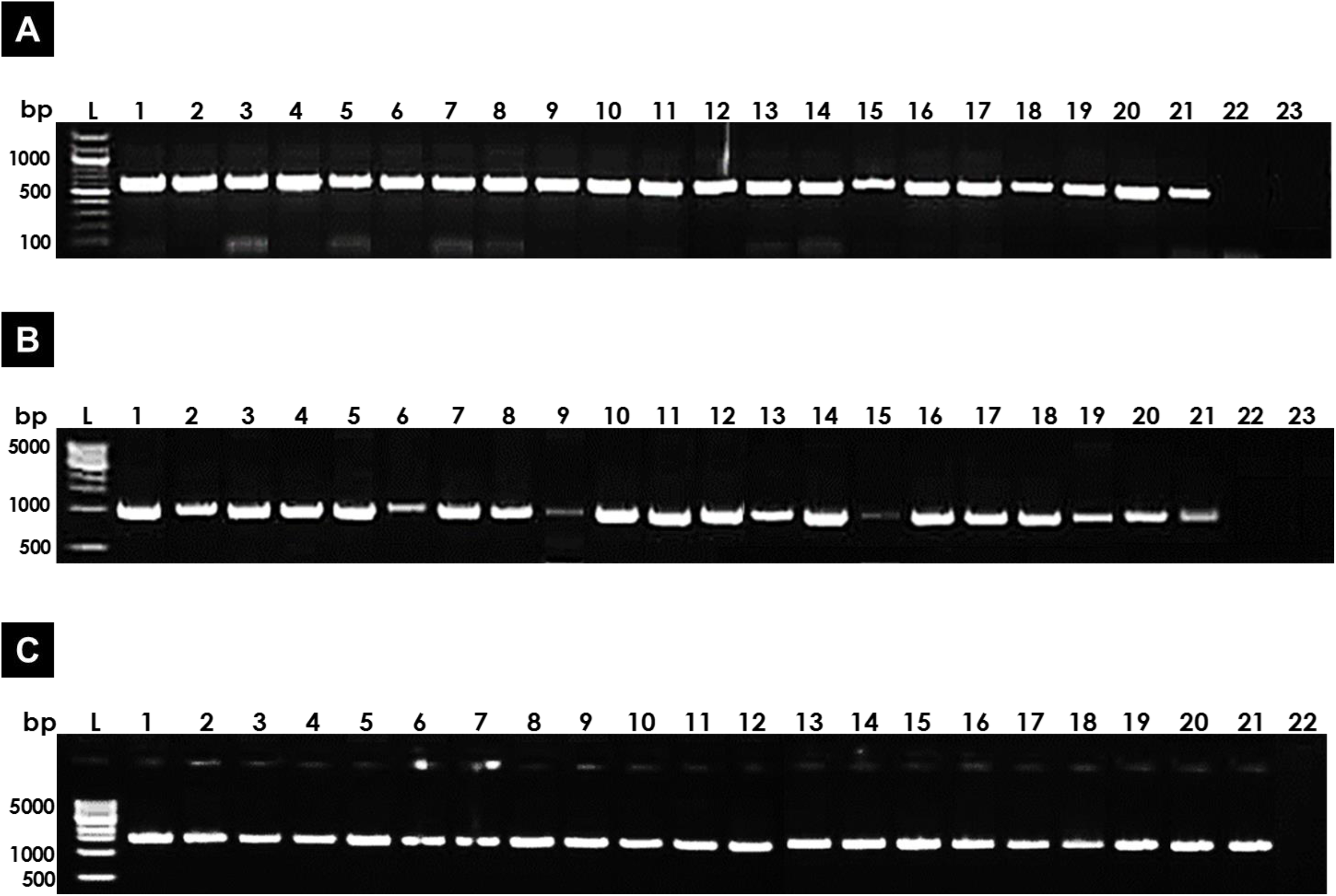
Molecular confirmation of plant-associated *P. aeruginosa.* (A) DNA amplification with *Pseudomonas* genus-specific primers PA-GS-F/R (amplicon size – 618 bp); Lane L, 100 bp marker. (B) PCR analysis with *P. aeruginosa* species-specific primers PA-SS-F/R (amplicon size – 956 bp); Lane L, 1 Kb marker; Lane 1 to 18, PPA01 to PPA18; Lane 19 to 21, ATCC10145, ATCC9027 and PAO1; Lane 22, *Enterobacter cloacae* (genera under gammaproteobacteria). Lane 23, negative control (no template). (C) PCR analysis with universal eubacterial primers (amplicon size – 1500 bp); Lane L, 1 Kb marker.

### Pyocyanin production

The PPA strains were qualitatively screened for their ability to produce pyocyanin (Fig. 2A). As controls, the three *P. aeruginosa* strains PAO1, ATCC10145, and ATCC9027 were included. In this assay, low levels of fluorescence were observed for PPA01, PPA06, PPA09, and PPA12. High level of fluorescence was observed for the remaining strains.

**Fig. 2.**
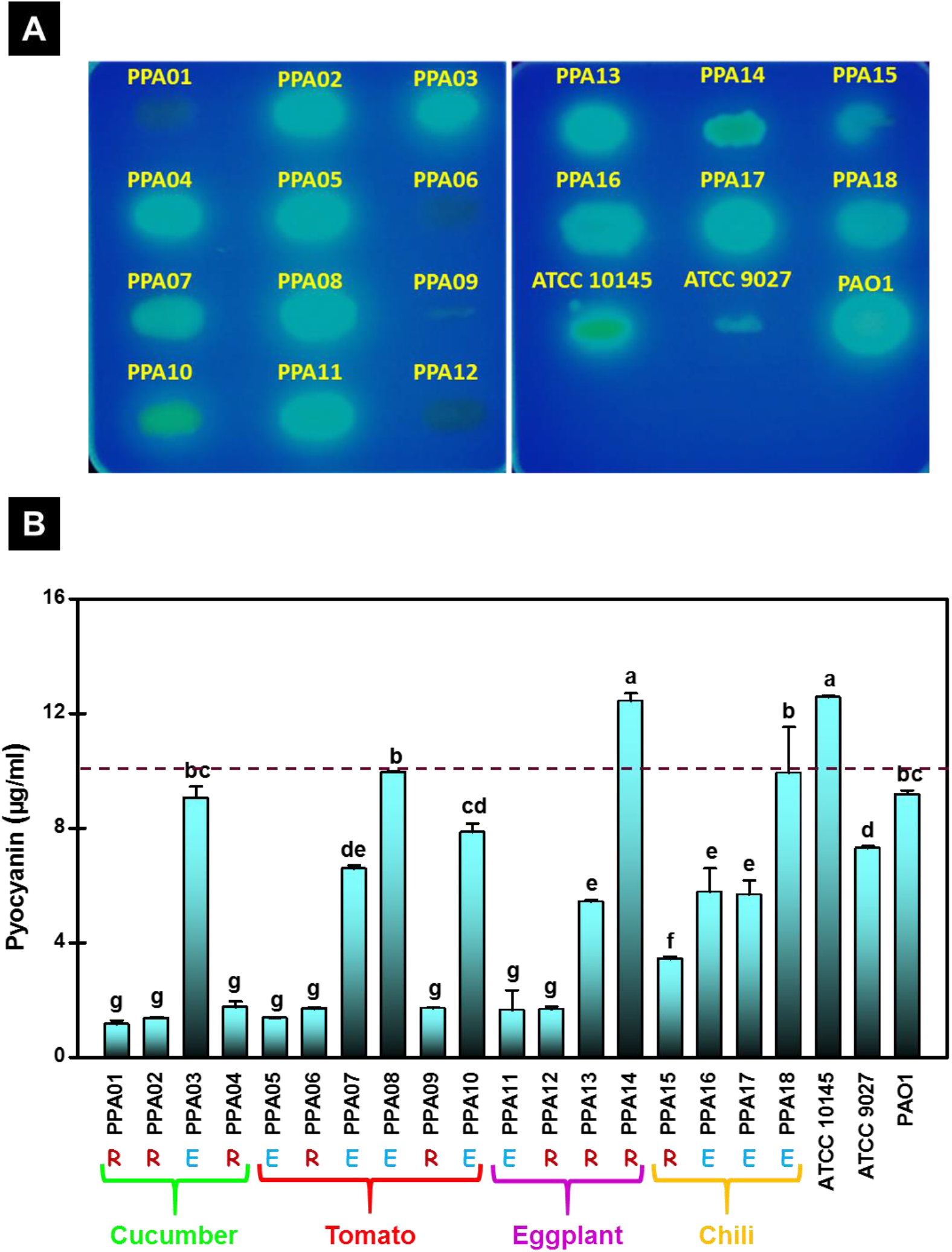
Pyocyanin production by *P. aeruginosa* strains. (A) *P. aeruginosa* strains grown on pyocyanin-specific medium are exhibiting fluorescence under ultra-violet radiation, indicative of pyocyanin. (B) Quantitative pyocyanin levels released by *P. aeruginosa* strains. Values plotted are mean of three replicates with standard errors and the alphabets above the bars indicate the ranking of strains (significant differences (p < 0.05) based on Duncan’s multiple range test (DMRT). The dashed line indicates the average levels of pyocyanin made by the clinical strains, ATCC10145, ATCC9027 and PAO1. R, rhizosphere strain; E, endophytic strain.

All the strains were grown in glycine-alanine broth to induce pyocyanin release (Ingledew and Campbell 1969). The pyocyanin was quantified and expressed as µg/ml (Fig. 2B). The significance of the values was analyzed using One-way ANOVA and DMRT (XLSTAT, version 2010.5.05). As expected, the control strains of *P. aeruginosa* released high levels of pyocyanin within 48 h of incubation (Fig. 2B). Except for one rhizospheric strain (PPA14), the rest produced significantly lower levels of pyocyanin. The pyocyanin produced by PPA14 is comparable to the control, ATCC10145 (designated by ‘a’). Except for one endophytic strain (PPA05), the rest produced pyocyanin comparable to the control strains PAO1 and ATCC9027. The four rhizospheric strains, PPA01, PPA06, PPA09, and PPA12, made low levels of pyocyanin (Fig. 2B) consistent with their low fluorescence seen in the plates (Fig. 2A). All PPA strains produced pyocyanin, indicating that they are indeed *P. aeruginosa*.

### Sequence alignment of full-length 16S rRNA genes

The complete length sequence (1500 bp) of 16SrDNA of the PPA strains were compared with the clinical strains (PAO1, ATCC10145, and ATCC9027) and previously identified plant-associated strains (from rice, guava, grass, pine, banana, lily, onion, ginseng and aloe vera) to determine their evolutionary relatedness. Sixteen of the PPA strains isolated in this study and nine of the agricultural strains from the previous studies had more than 97% sequence identity with PAO1 (Fig. 3). PPAO8, and PPA15, isolated from tomato, and chili in this study, and SEGB6, isolated from guava in previous study (Accession no: MN565979) showed less than 97% identity with PAO1. However, PPA08, and PPA15 were included for further analysis as they got amplified by the species-specific primers (Fig. 1B) and could produce pyocyanin (Fig. 2), the biomarker of *P. aeruginosa*.

**Fig. 3.**
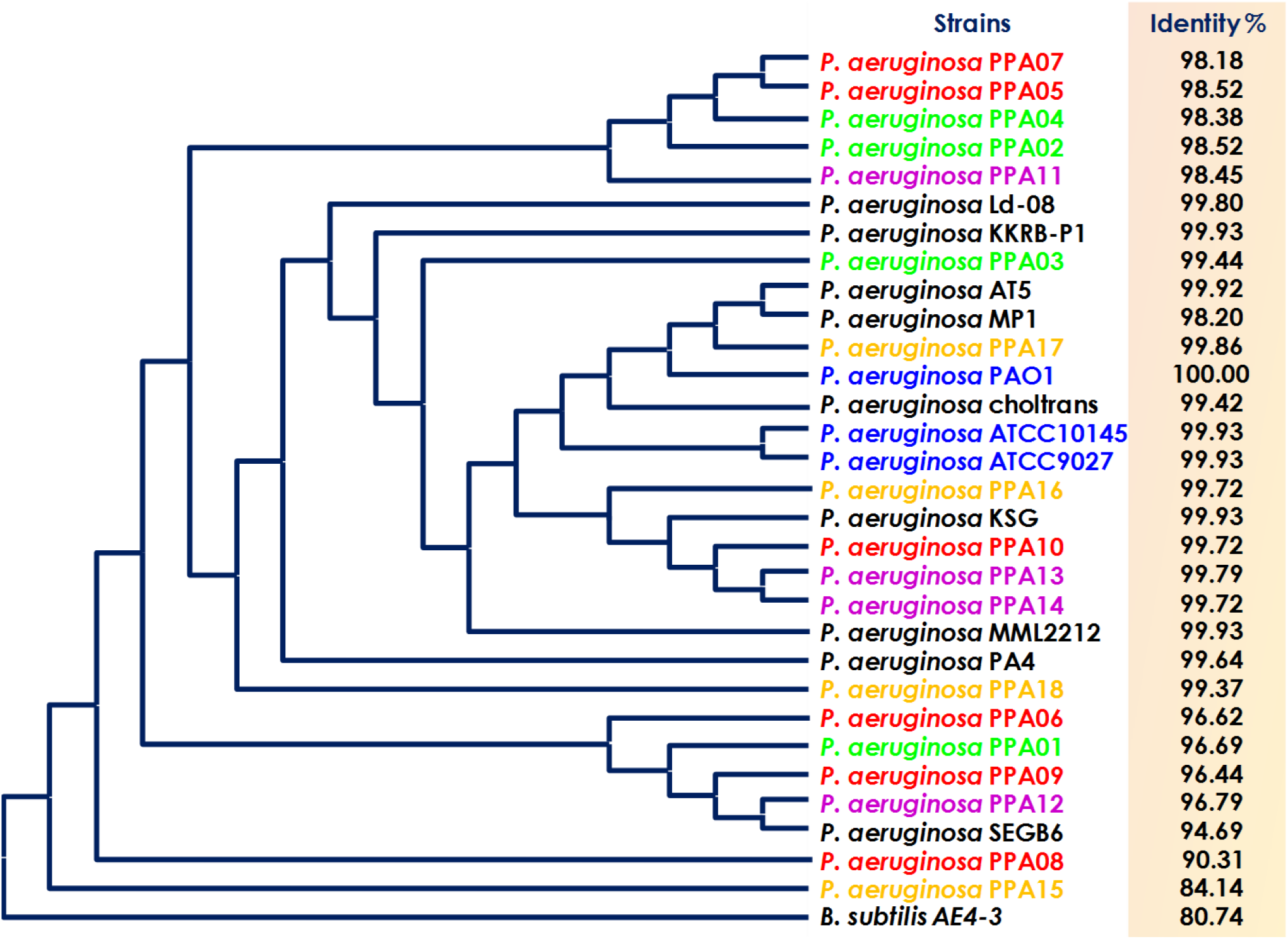
Phylogenetic and homology analyses of *P. aeruginosa* strains. Phylogenetic tree based on Neighbor-Joining method constructed with 16S rDNA sequence of *P. aeruginosa* strains isolated in the current study (PPA01 to PPA18) from cucumber (green font), tomato (red font), eggplant (purple font), and chili (yellow plant); The clinical strains used are ATCC10145, ATCC9027, and PAO1 (blue font). Previously characterized agricultural *P. aeruginosa* strains (black font) are from various niches: *P. aeruginosa* AT5 (Accession No: MN636767) and *P. aeruginosa* PA4 (Accession No: MN636761), ginseng leaf, and rhizoplane, respectively. *P. aeruginosa* SEGB6 (Accession No: MN565979) is from guava leaf; and *P. aeruginosa* Ld-08 16S (Accession No: MT472133) is a lily endophyte. The following are rhizosphere isolates: *P. aeruginosa* KSG (Accession No: LN874213, grass); *P. aeruginosa* MML2212 (Accession No: EU344794, rice); *P. aeruginosa* KKRB-P1 (Accession No: MW149279, pine); *P. aeruginosa* MP1 (Accession No: MT937234, banana); *P. aeruginosa* VL4 (Accession No: MN611376, onion), and *P. aeruginosa* choltrans (Accession No: MK782058, aloe vera). The 16S rDNA sequence of each strain was subjected to pairwise BLAST against PAO1.

*B. subtilis* was used as the outlier to create a phylogenetic tree using the Neighbor-Joining method (Fig. 3; Saitou and Nei (1987)) to study the evolutionary relatedness of the strains. Evolutionary distances of the strains were computed using the Maximum Composite Likelihood method (Tamura et al. 2004). The plant-associated strains clustered together with the clinical strains indicating their evolutionary relatedness (Fig. 3). Three pairs of *P. aeruginosa* strains had 99% sequence identity and co-clustered; clinical strains (ATCC10145, and ATCC9027), eggplant rhizosphere strains (PPA13, and PPA14), and tomato endophytes (PPA05, and PPA07). *P. aeruginosa* strain AT5 from ginseng leaf, and strain MP1 from banana rhizosphere co-clustered regardless of their niches. In addition, one of the eggplant rhizosphere strain, PPA12 isolated in the current study clustered together with a previously identified guava leaf isolate. Phylogenetic analyses showed that the *P. aeruginosa* strains from different niches could have high relatedness based on their 16s DNA sequence.

### Molecular typing to determine genetic diversity

The 18 *P. aeruginosa* isolates (PPA01 to PPA18) were fingerprinted using ERIC and BOX primers to determine their clonal diversity (Versalovic et al. 1994). The ERIC sequences are the conserved palindromic regions (126 bp) present in multiple copies within a bacterial genome (Wilson and Sharp 2006). The BOX sequences are repetitive elements comprised of three areas, *boxA* (59 bp), *boxB* (45 bp), and *boxC* (50 bp) (Versalovic et al. 1994). The location of ERIC and BOX regions differ between the strains. These variations are exploited to reveal the strain-level genetic heterogeneity (Wilson and Sharp 2006).

The plant-associated and control *P. aeruginosa* strains (PAO1, ATCC10145, and ATCC9027) were amplified (Fig. 4A and B) using the ERIC (ERIC 2F/IR; Fig. 4A) and the BOX (BOXA1; Fig. 4B) primers (Table 3). None of the PPA strains had a similar fingerprint to the control strains (Lanes 19 to 21; Fig. 4A and B). Some of the PPA strains (PPA02, PPA04, PPA05, and PPA07) had visually similar fingerprints.

**Fig. 4.**
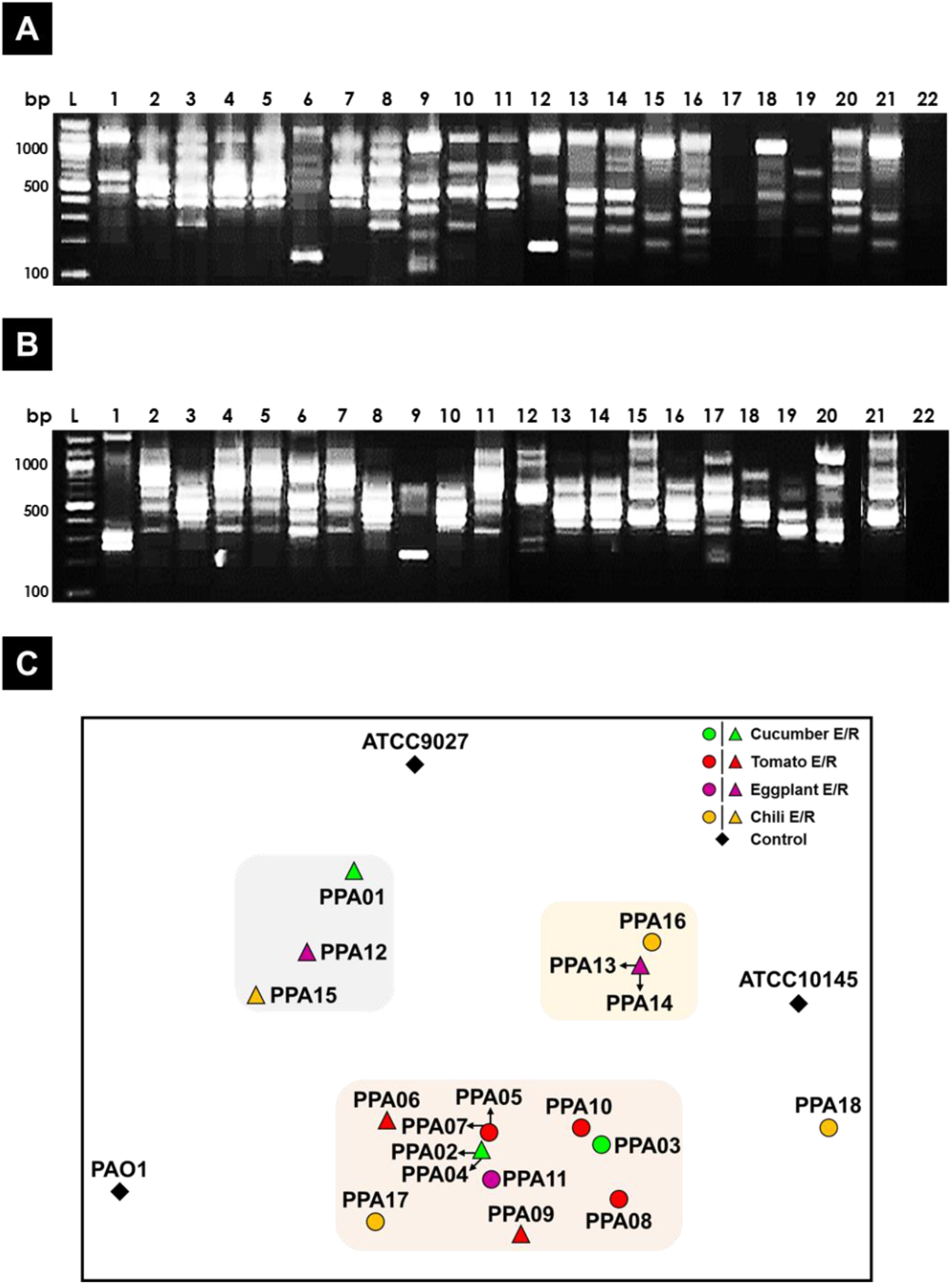
DNA fingerprinting profile of *P. aeruginosa* strains. Fingerprinting was done as described in materials and methods (A) ERIC fingerprint using ERIC2F and ERIC1R primers (Table 2). (B) BOX fingerprint using BOXA1 primer (Table 2). Lane 1 to 18, PPAO1 to PPA18; Lane 19 to 21, ATCC10145, ATCC9027 and PAO1; Lane 22, negative control (no template). (C) Bray-Curtis similarity-based non-metric multi-dimensional scaling (MDS) of the DNA fingerprinting pattern. The strains are color-coded based on their plant source, cucumber (green), tomato (red), eggplant (purple), and chili (yellow). R, rhizosphere strain; E, endophytic strain.

The fingerprint data were analyzed using multidimensional scaling (MDS) based on the Bray-Curtis similarity matrix (Clarke 1993) to determine the similarity between the strains (Fig. 4C). All three control strains, PAO1, ATCC10145, and ATCC9027, were scattered away from the plant-associated strains (Fig. 4C). The PPA strains clustered into three groups. Cluster A was made of three rhizospheric strains, PPA01, PPA12, and PPA15, from three different crops, cucumber, eggplant, and chili, respectively (Fig. 4C). Cluster B had a chili endophyte (PPA16) and two eggplant rhizospheric strains (PPA13 and PPA14). Cluster C had 11 PPA strains isolated from all four plants. All the rhizospheric and endophytic strains from the tomato plant co-clustered in one group (cluster 3). PPA18, the chili endophyte, had a unique fingerprint and did not cluster with any other strains. Fingerprinting data showed that the strains except for the tomato isolates did not cluster by their source (rhizosphere or endophyte). The tomato strains are more clonal than the others.

### Ability to solubilize complex soil minerals

Plant growth-promoting rhizobacteria secrete organic acids to solubilize complex soil-bound minerals and release nutrients such as phosphorous, potassium, and zinc (Oleńska et al. 2020). The host plants absorb these nutrients for their growth and development. Thus, we postulated that the *P. aeruginosa* strains in an agricultural setting might promote plant growth by solubilizing complex soil minerals. The ability of the PPA strains to release phosphorous, potassium, and zinc was tested (Fig. 5). The three control isolates of human origin used in this study, PAO1, ATCC10145, and ATCC9027, were previously never tested for their ability to solubilize minerals.

**Fig. 5.**
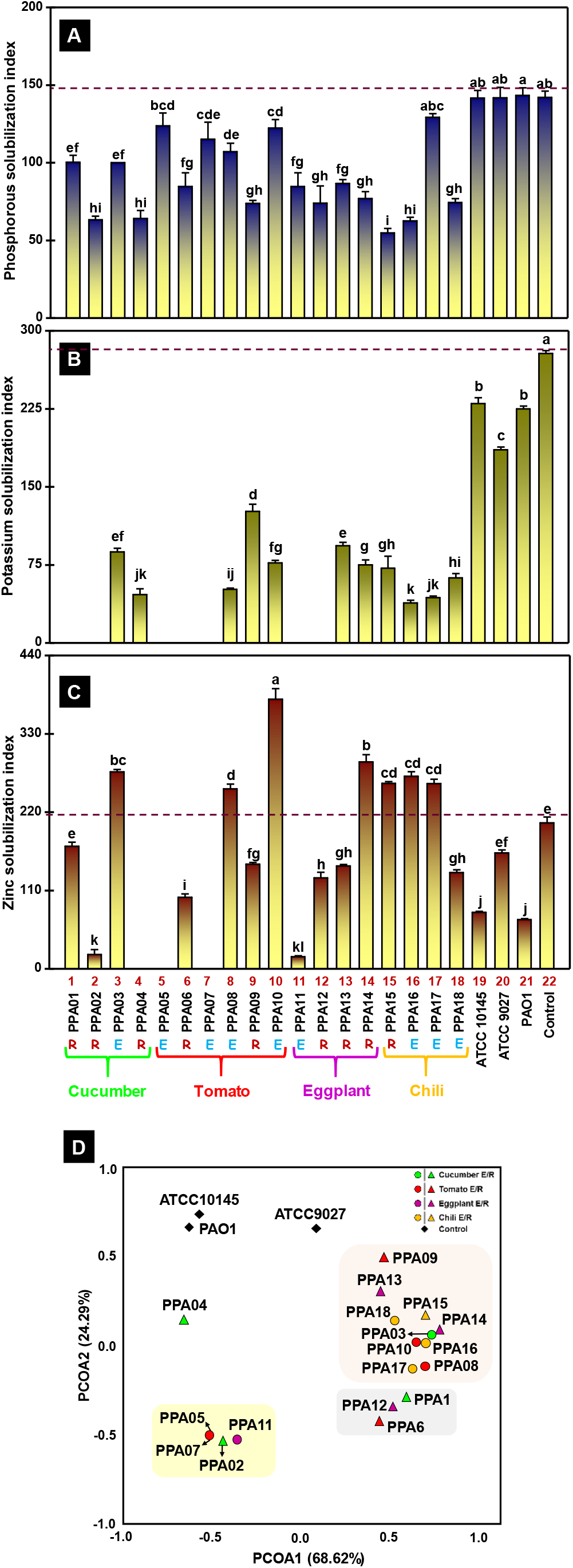
Mineral solubilization ability of *P. aeruginosa* strains. The graphs represent the phosphorous (A), potassium (B), and zinc (C) solubilization index of *P. aeruginosa* strains. Values plotted are the mean of three replicates with standard errors. The alphabets above the bars indicate the ranking of strains (significant differences (*p* < 0.05) based on Duncan’s multiple range test (DMRT). The dashed line indicates the level made by the control strains, *Bacillus subtilis* var. *phospaticum* strain Pb1 (Phosphorous-solubilizing bacteria), *Paenibacillus mucilaginosus* strain KRB9 (Potassium-releasing bacteria), and *Pseudomonas chloraraphis* strains ZSB15 (Zinc-solubilizing bacteria). The strains are color-coded based on their plant source, cucumber (green), tomato (red), eggplant (purple), and chili (yellow). R, rhizosphere strains; E, endophytic strain. (D) Principal coordinate analysis (PCoA) based on Euclidean distance on mineral solubilization potential of *P. aeruginosa* strains. The percentage values in parentheses x- (PCoA1) and y-axes (PCoA2) depict the similarities and deviations among the strains based on their mineral solubilizing abilities.

#### Phosphorous release

The ability of *P. aeruginosa* strains (controls and PPAs) to release phosphorous from tri-calcium phosphate (insoluble mineral complex) was tested and presented as phosphorous solubilization index (PSI, Fig. 5A). *B. megaterium* var *phosphaticum* Pb1, a well-characterized phosphate solubilizing bacteria, was used as the positive control (Balamurugan and Gunasekaran 1996; Gomathy et al. 2007). As expected, obtained a high PSI value for the control (Bar 22; Fig. 5A). Human-associated *P. aeruginosa* strains, PAO1, ATCC10145, and ATCC9027, had a phosphorous solubilization index (PSI) similar to the control (Bars 19-21, respectively; Fig. 5A). All the PPA strains solubilized tri-calcium phosphate at varying levels. One-way analysis of variance (ANOVA) and Duncan’s multiple range test (DMRT) demonstrated that some of the PPA strains (Bars 2, 4, 15, and 16) had significantly lower PSI values. All the four tomato endophytes, PPA05, PPA07, PPA08, and PPA10, had comparably high PSI values as indicated by the shared letter ‘d’ (Fig. 5A). The PPA15 (chili rhizosphere) and PPA17 (chili endophyte) strains had the minimum (Bar 15) and maximum (Bar 17) PSI values, respectively. Though this analysis only includes 18 strains, it appears that the top five PPA strains (PPA05, PPA07, PPA08, and PPA17) that had a high ability to release phosphorous were from the endophytic niche.

#### Potassium release

*P. aeruginosa* strains were tested for their ability to solubilize potassium aluminum silicate, and presented as potassium solubilization index (KSI, Fig. 5B). As a control, well-characterized potassium releasing bacteria, *B. mucilaginous* KRB9 was used (Brindavathy and Gopalaswamy 2017). As expected, the control strain had a high KSI value (Bar 22; Fig. 5B). One-way ANOVA and DMRT demonstrated that the KSI value of the control strain was significantly higher (indicated by letter ‘a’) than the *P. aeruginosa* strains. All the chili isolates had the ability to solubilize potassium aluminium silicate (Bars 15-18). Three endophytic PPA strains (Bar 5, 7, and 11) from tomato and eggplant and four rhizospheric PPA strains (Bar 1, 2, 6, and 12) from cucumber, tomato, and eggplant could not release potassium from the mineral complex. More importantly, the KSI of the tested human-associated *P. aeruginosa* isolates (Bar 19-21) was higher than that of PPA strains.

#### Zinc release

The zinc solubilization (ZS) potential of the *P. aeruginosa* strains was tested and presented as an index (ZSI, Fig. 5C). A well-characterized zinc-solubilizing bacterium *P. chlororaphis* ZSB15 was used as a control (Bowya and Balachandar 2020). Interestingly, the control strain had significantly lower ZSI (Bar 22) than 39% of the PPA strains (Bar 3, 8, 10, 14, 15, 16, and 17). The ZSI of the tomato endophyte PPA10 (Bar 10) was significantly higher than the control (Bar 22; Fig. 5C). Two tomato endophytes, (PPA05 and PPA07,) and the cucumber rhizosphere strain PPA04 could not solubilize zinc oxide. The human-associated *P. aeruginosa* strains had very low ZSI values (Bar 19-21).

#### Clustering based on mineral solubilization index

Euclidean distance-based principal coordinate analysis (PCoA) (NCSS, Kaysville, USA) determined the similarity of *P. aeruginosa* strains based on their ability to solubilize the complex soil minerals. The *P. aeruginosa* controls (PAO1, ATCC10145, and ATCC9027) clustered away from the PPA strains (Fig. 5D). All the PPA strains were grouped into three clusters except for a cucumber rhizosphere strain, PPA04. Cluster A was occupied by two tomato endophytes (PPA05 and PPA07), an eggplant endophyte (PPA11), and a cucumber rhizosphere strain (PPA02). All the chili isolates (PPA15-PPA18) grouped in cluster B along with five other strains from cucumber, tomato, and eggplant. Cluster C had three endophytic strains from cucumber (PPA01), tomato (PPA06), and eggplant (PPA12). In the PCOA plot based on mineral solubilization properties, the PPA strains except for the chili isolates did not cluster by their plant source or niche (rhizosphere or endophyte).

### Ammonification potential

Terrestrial plants can neither access the gaseous form of nitrogen from the atmosphere nor the organic form of nitrogen from the soil. The plants solely depend on associated microbes to release ammonia (ammonification) and nitrate (nitrification) through the decomposition of soil organic matter (Oleńska et al. 2020). Thus, we hypothesized that the agricultural strains of *P. aeruginosa* might contribute to host plant growth through the nitrogen cycle.

Tested the ability of the *P. aeruginosa* strains to convert nitrogen into ammonia by quantifying the amount of ammonia released into peptone broth (Cappucino and Sherman, 1992). *B. altitudinis* FD48, a well characterized plant growth-promoting bacteria, was used as a control (Kumar et al. 2017; Ambreetha et al. 2018; Narayanasamy et al. 2020). All the strains can convert nitrogen to ammonia. The amount of ammonia released by the control (Fig. 6, FD48) was significantly lower than most of the *P. aeruginosa* strains (62%). The endophytes from cucumber (PPA03), and tomato (PPA08) had significantly higher ammonification activity as per ANOVA and DMRT analyses (indicated by ‘a’). The top four ammonifiers in the PPA group (PPA03, PPA08, PPA10, and PPA18) were all from endophytic niches. Least ammonification was observed with two rhizospheric strains (PPA01, and PPA06 from cucumber, and tomato, respectively), and an eggplant endophyte, PPA11 (indicated by ‘j’, Fig. 6). *P. aeruginosa* ATCC10145 produced similar ammonia levels compared to the control strain FD48. However, *P. aeruginosa* ATCC9027 and PAO1 produced significantly higher ammonia levels. It appears that the ammonification potential is not dependent on the source of isolation. However, 63% of the strains that had high ammonification activity were endophytes.

**Fig. 6.**
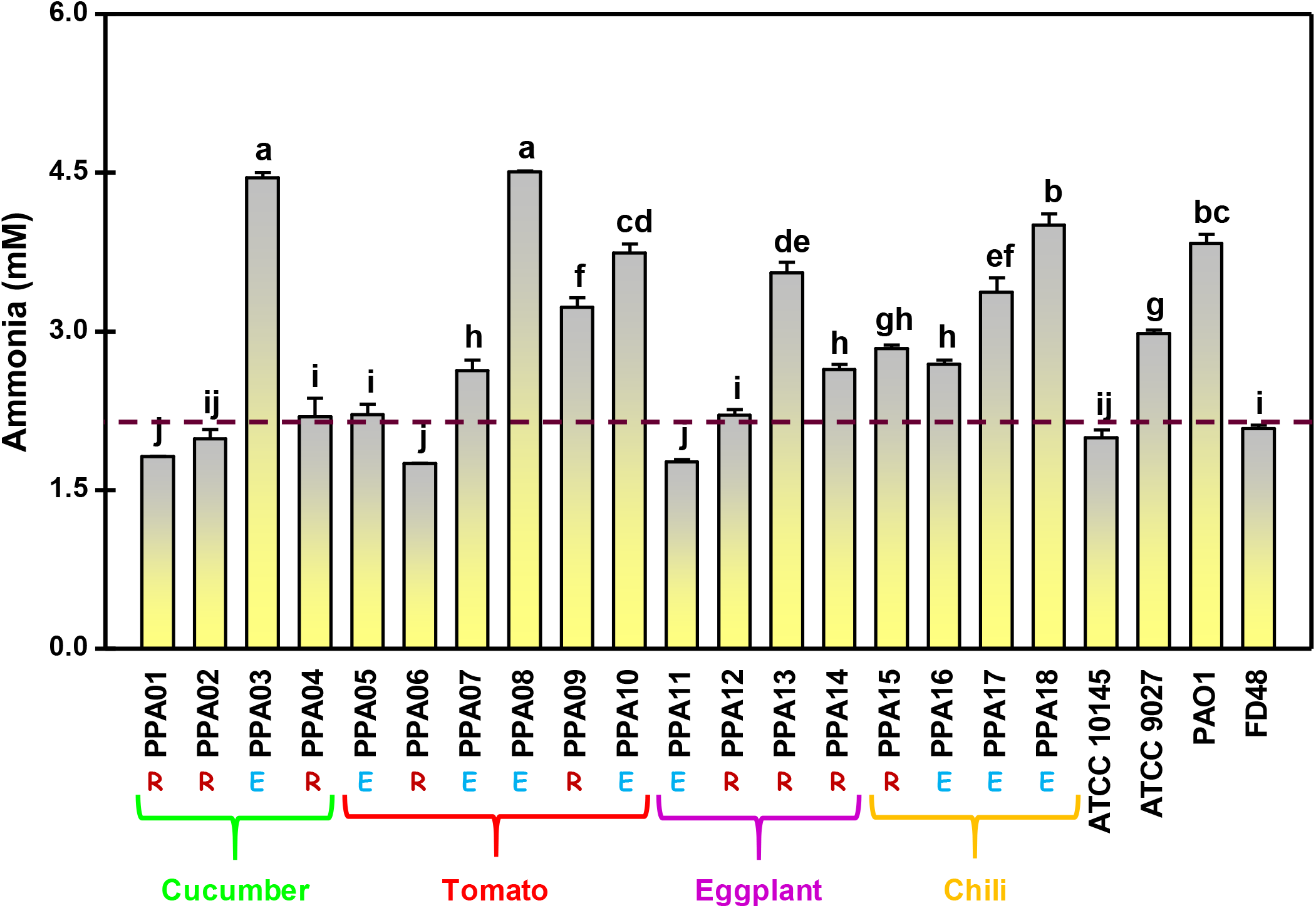
Ammonia production by *P. aeruginosa* strains. The graph represents the ammonia (mM) released by *P. aeruginosa*. Values plotted are the mean of three replicates with standard errors, and the alphabets above the bars indicate the ranking of strains (significant differences (*p* < 0.05) based on Duncan’s multiple range test (DMRT). The dashed line indicates the level produced by the control, *Bacillus altitudinis* strain FD48. The strains are color-coded based on their plant source, cucumber (green), tomato (red), eggplant (purple), and chili (yellow). R, rhizosphere strain; E, endophytic strain.

### Ability to release siderophores

Iron is an essential element required by all living organisms, but its bioavailability is highly limited in the soil (Colombo et al. 2014). Microbes that could release siderophores scavenge the iron molecules from the soil leading to iron starvation and death of other competing counterparts (Leong 1986). Siderophore-producing microbes act as biocontrol agents by inhibiting the growth of soil-borne pathogens (Ghosh et al. 2020). Microbial siderophores also induce plant immunity making their host plant disease-resistant (Aznar et al. 2014). In this study, it was hypothesized that plant-associated *P. aeruginosa* confers protection through siderophore production.

The ability of *P. aeruginosa* strains to release siderophores was qualitatively tested in chrome azurol S (CAS) agar medium (Schwyn and Neilands 1987). Hexadecyltrimethylammonium bromide, and iron(III) form a blue-colored dye-iron complex in CAS-containing agar medium. Siderophores from *P. aeruginosa* scavenges the iron(III) from this complex leading to yellow coloration. The formation of yellow zones confirmed the release of siderophores by all the tested strains (Fig. 7A). The amount of siderophore released was then quantified (Fig. 7B). The plant growth-promoting strain *B. altitudinis* FD48 was used as the positive control (Kumar et al. 2017; Ambreetha et al. 2018; Narayanasamy et al. 2020). As expected, the control and all the strains produced siderophores (Fig. 7B). ANOVA and DMRT analyses determined the significant level of variations in siderophores released by the tested strains (strains that shared alphabets had no significant difference with each other, but significantly higher than the control strain). The levels of siderophore produced in most of the plant-associated *P. aeruginosa* strains (63%) were higher than the control. The tomato rhizosphere strain PPA09 hyperproduced siderophore (Indicated by ‘a’). Similar to the plant isolates, there is a variation between human isolates in their ability to produce siderophores, PAO1 and ATCC10145 producing higher than the control. Both rhizospheric and endophytic *P. aeruginosa* isolates had the ability to release siderophores which likely benefits their host plants.

**Fig. 7.**
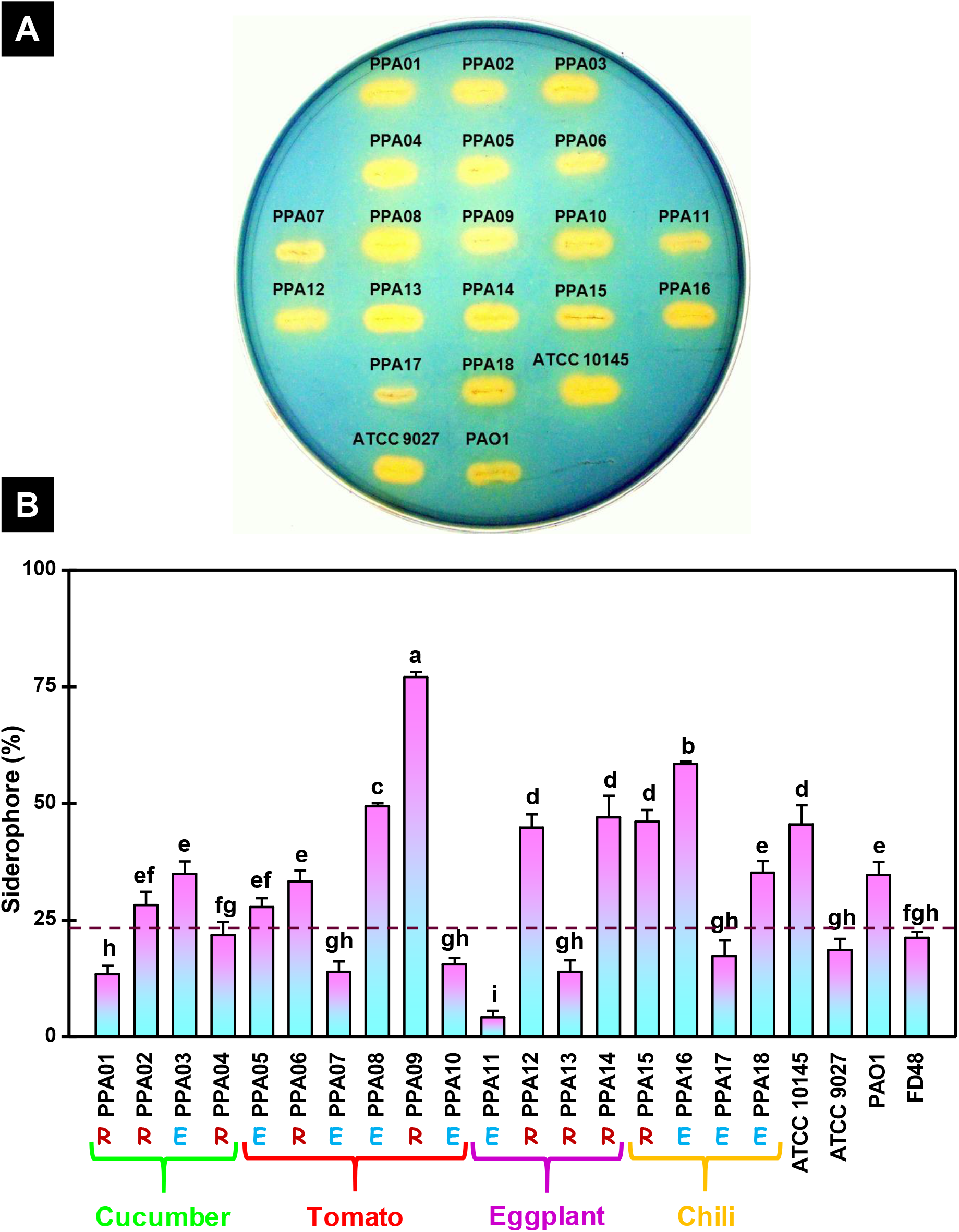
Siderophore production by *P. aeruginosa* strains. (A) Qualitative detection of siderophores based on the presence of yellow zones around the *P. aeruginosa* strains on chrome azurol S (CAS) agar plate (blue). (B) Quantitative analysis of siderophores released by *P. aeruginosa*. Values plotted are the mean of three replicates with standard errors, and the alphabets above the bars indicate the ranking of strains (significant differences (*p* < 0.05) based on Duncan’s multiple range test (DMRT). The dashed line indicates the level produced by the control, *Bacillus altitudinis* strain FD48. The strains are color-coded based on their plant source, cucumber (green), tomato (red), eggplant (purple), and chili (yellow). R, rhizosphere strain; E, endophytic strain.

### Indole acetic acid production

The natural auxin, indole acetic acid (IAA) is a phytohormone that plays a vital role in plant growth (Zhao 2010). Plants root exudates contain tryptophan that can be converted into IAA by the associated microbes (Kravchenko et al. 2004). Exogenous auxin released by the rhizospheric and endophytic bacteria contributes to cell division, cell elongation, improvement of the root architecture, and development of leaves, fruits, and flowers in the host plant (Sukumar et al. 2013; Ali et al. 2017; Ambreetha and Balachandar 2019). In other words, the plant-associated bacteria that can release auxin are considered beneficial to their host plants.

The ability of the plant-associated *P. aerugionsa* to convert exogenous tryptophan into IAA was investigated using Salkwoski’s reagent (Gordon and Weber 1951) and expressed as parts per million (ppm; Fig. 8). The statistical significance in the amount of IAA released was determined using the ANOVA and DMRT (strains that shared alphabets had no significant difference, Fig. 8). All strains, including the control *B. altitudinis* FD48, produced IAA. *P. aeruginosa* strains of non-plant origin had relatively lower levels of IAA than the control strain. Nearly 60% of the plant-associated strains released a higher amount of 1AA than the control (PPA 02, 04, 05, 06, 07, 08, 09, 11, 15, 16, and 18). The tomato rhizospheric strain PPA09 had the highest IAA levels, whereas eggplant-rhizosphere strains (PPA12, PPA13, and PPA14) had the lowest. Two rhizospheric cucumber strains, PPA02, and PPA04 had similar IAA production (indicated by ‘b’). Endophytes from eggplant (PPA11) and chili (PPA16) released similar IAA levels (indicated by ‘e’). In summary, both endophytic and rhizospheric *P. aeruginosa* isolates are capable of releasing extracellular IAA from tryptophan.

**Fig. 8.**
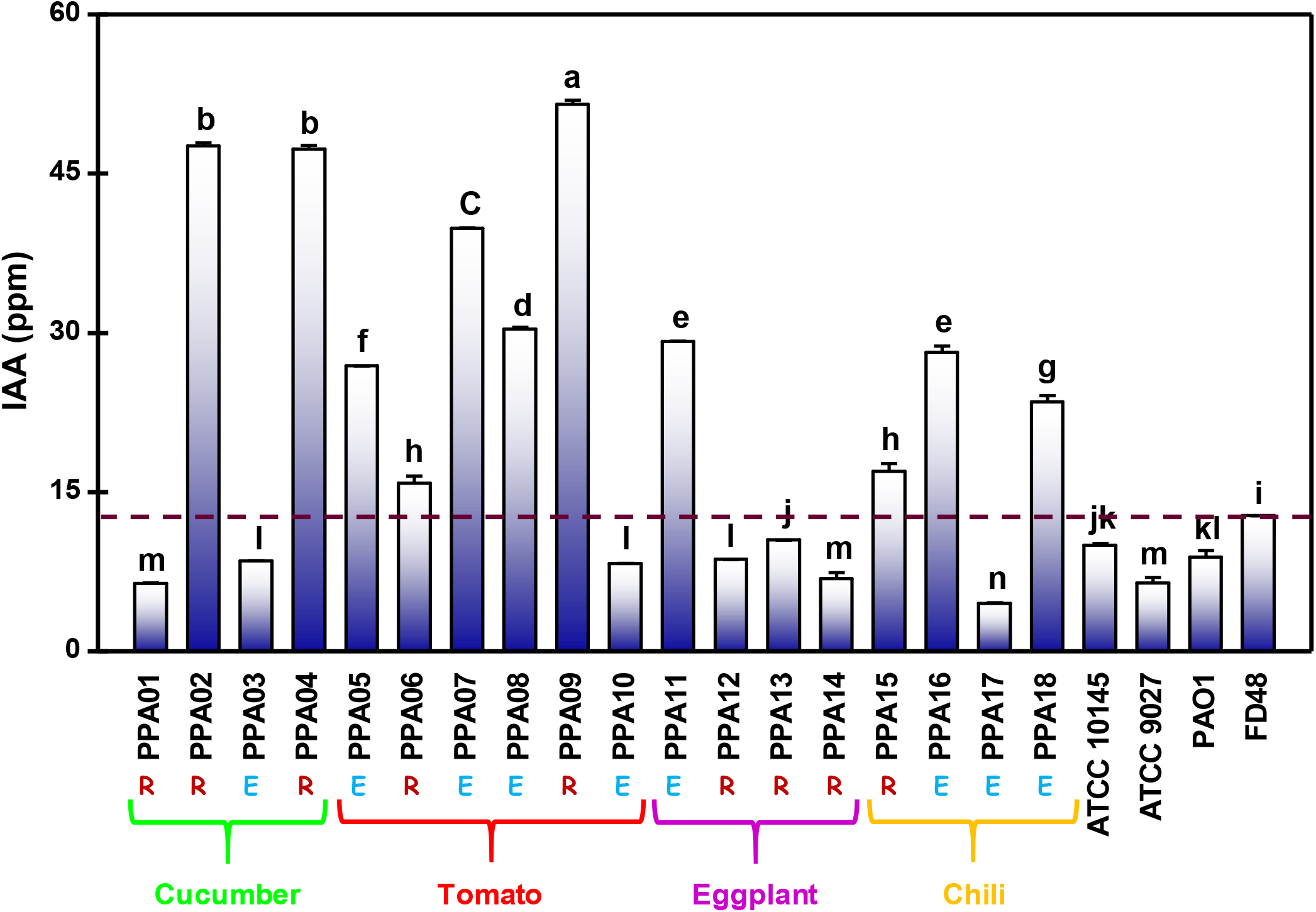
Indole acetic acid (IAA) production by *P. aeruginosa* strains. The graph represents IAA released by *P. aeruginosa* in parts per million (ppm). Values plotted are the mean of three replicates with standard errors, and the alphabets above the bars indicate the ranking of strains (significant differences (p < 0.05) based on Duncan’s multiple range test (DMRT). The dashed line indicates the level produced by the control, *Bacillus altitudinis* strain FD48. The strains are color-coded based on their plant source, cucumber (green), tomato (red), eggplant (purple), and chili (yellow). R, rhizosphere strain; E, endophytic strain.

## Discussion

Pseudomonads is the most abundant bacterial community in the soil (Janssen 2006) and in some areas nearly 78% of the Pseudomonads is *P. aeruginosa* (Noura et al. 2009). The initial assumption that *P. aeruginosa* inhabiting plants and humans are two different species was defenestrated by demonstrating the ability of clinical *P. aeruginosa* to colonize plants (Schroth et al. 1977; Schroth et al. 2018). Further, Alonso et al. (1999) have demonstrated that environmental isolates from oil-contaminated soil harbor virulence and antibiotic resistance similar to the clinical *P. aeruginosa* strains. However, there are limited molecular and biochemical evidence to prove the relatedness of human- and plant-associated *P. aeruginosa* strains. The current study reveals the shared evolutionary and functional attributes of clinical and agricultural isolates of *P. aeruginosa*. In addition, we also found that the clinical isolates tested here harbor plant growth-promoting capabilities.

### *P. aeruginosa* found in rhizosphere and internal tissues of vegetable plants

Seven different edible plants (Table 3) grown in Southern India were tested for the presence of *P. aeruginosa.* Genus- and species-specific primers (Fig. 1) previously developed for clinical *P. aeruginosa* were used for molecular identification of the agricultural strains (Spilker et al. 2004). Eighteen different strains of *P. aeruginosa* were identified in rhizosphere and endophytic niches of cucumber, tomato, chili, and eggplant (Table 1). We could not detect *P. aeruginosa* in rice, which might be due to technical and sampling limitations. However, there are previous reports on the presence of *P. aeruginosa* in rice ecosystem (Shanmugaiah et al. 2010; Shi et al. 2015).

There have been regular reports on *P. aeruginosa* contamination in vegetables at supermarkets, hospital kitchens, canteens, and street vendors (Kominos et al. 1972; Wright et al. 1976; Correa et al. 1991; Viswanathan and Kaur 2001; Allydice-Francis and Brown 2012; Nithya and Babu 2017). Such contaminations could have occurred through different exposures during handling, processing, packing, transportation, or storage. A few *P. aeruginosa* strains have been previously isolated and characterized from chili (Linu et al. 2019) pepper (Kumar et al. 2013), tomato (Iasur Kruh et al. 2020), medicinal plant *Achyranthes aspera* L. (Devi et al. 2017), ginger (Jasim et al. 2014), reed (Wu et al. 2018), chickpea (Mukherjee et al. 2020), aloe vera (Akinsanya et al. 2015), sugar cane (Singh et al. 2021), and wheat (Sun et al. 2021). In this study, we have found *P. aeruginosa* in rhizosperic and endophytic association with the vegetable plants harvested directly from the farm. In any case, the presence of human pathogens in fresh vegetables or their plants indicate the potential health hazard associated with agricultural produces (Berger et al. 2010; Al-Kharousi et al. 2016).

### *P. aeruginosa* can promote plant growth

*P. aeruginosa* strains tested in this investigation had numerous plant growth-promoting attributes. They had the ability to solubilize complex soil minerals and release available form of nutrients (Fig. 5). The clinical strains could also solubilize all three mineral complexes (tri-calcium phosphate, potassium aluminium silicate, and zinc oxide) (Fig. 5). Nearly 61% and 83% of the plant-associated strains released potassium and zinc, respectively, whereas all the PPA strains could release phosphorous from the tri-calcium phosphate (Fig. 5). *P. aeruginosa* strains with mineral solubilizing ability are known to improve the growth and productivity of vegetable crops such as green gram, tomato, okra, and African spinach (Adesemoye et al. 2008; Ahemad and Khan 2010). Recently, use of mineral weathering bacteria is recommended to convert rocks and minerals into potential plant fertilizers (Ribeiro et al. 2020).

All the strains tested in this study had the ability to release siderophores and IAA which is likely to benefit the associated plants. Previous studies have demonstrated that siderophores and IAA released by *P. aeruginosa* improves seed germination, root length and shoot length, of the host plants (Sulochana et al. 2014; Sah et al. 2017). *P. aeruginosa* siderophores have been proven as effective inhibitors of plant pathogens such as *Fusarium oxysporum, Trichoderma herizum, Alternaria alternate,* and *Macrophomina phasiolina* (Bano and Musarrat 2003). Nearly 50% of the strains tested in this study released excessive levels of siderophores and IAA (Fig.7B and 8), which directly helps in plant growth-promotion and disease protection (Hariprasad et al. 2014; Marathe et al. 2017).

Pyocyanin, a redox-active phenazine compound was released by all the *P. aeruginosa* strains tested in this study (Fig. 2B). Pyocyanin accounts for 90% of this bacterium’s biocontrol ability (Waksman and Woodruff 1940; Abou Raji El Feghali and Nawas 2018). It protects the host plants by inhibiting the growth of other soil pathogens (Anjaiah et al. 2003; Mahmoud et al. 2016). In addition, pyocyanin triggers induced systemic resistance of the host plant against various fungal pathogens (Audenaert et al. 2002; De Vleesschauwer et al. 2006). In the current study, 77% of the endophytic strains and 33% of rhizospheric strains had high levels of pyocyanin (Fig. 2B). In eukaryotic organisms, pyocyanin is cytotoxic to the respiratory, urological, central nervous, and vascular systems (Hall et al. 2016). However, the pyocyanin dosage required for plant protection is non-lethal to the eukaryotic cells (Priyaja et al. 2016). As most of the PPA strains had high pyocyanin and siderophore levels, their ability to inhibit bacterial and fungal phytopathogens will be tested in the future.

The three control isolates of human origin used in this study, PAO1, ATCC10145, and ATCC9027, were previously never tested for their ability to promote plant growth. These strains had the plant-beneficial traits comparable to the agricultural *P. aeruginosa* (Fig. 5-8). This further argues for the evolutionary relatedness of the plant- and human-associated strains.

### Conflicting reports on agricultural *P. aeruginosa*

Previous studies have given conflicting reports on the interactions between *P. aeruginosa* and its host plant. Several studies have demonstrated that *P. aeruginosa* causes rot and wilt (Clara 1930; Elrod and Braun 1942; Cother et al. 1976; El-Said et al. 1982; Bradbury 1986; Gupta et al. 1986; Gao et al. 2014; Tiwari and Singh 2017). On the other hand, plant-beneficial properties of agricultural *P. aeruginosa,* including a reduction in plant disease incidence have been reported in multiple investigations (Ali Siddiqui and Ehteshamul-Haque 2001; Adesemoye and Ugoji 2009; Yasmin et al. 2014; Radhapriya et al. 2015; Arif et al. 2016; Durairaj et al. 2017; Gupta and Buch 2019; Chandra et al. 2020).

Though, the two clinical strains, PAO1 and ATCC10145 used in the current study were initially isolated from burn wound and outer ear infection (Holloway 1955; Picard et al. 1994) they possessed plant-beneficial characteristics (Fig. 5-8). Likewise, there is a possibility for agricultural strains to harbor human virulence factors. Kumar et al. (2013) demonstrated that endophytic *P. aeruginosa* strains isolated from black pepper protected their host from fungus and nematode but they were cytotoxic to mammalian A549 cells, analogous to clinical strains. In 1980’s it was demonstrated that both plant and human strains of *P. aeruginosa* could infect animals (Lebeda et al. 1984). The clinical *P. aeruginosa* strain, PA14 harbors conserved virulence factors that could elicit disease in both plant and animal system (Schroth et al. 1977; Rahme et al. 2000; He et al. 2004). In lieu of these findings, the use of *P. aeruginosa* in agricultural setting is not recommended. Despite their agricultural benefits, this bacterium could be a potential health hazard to farm workers, farm animals, and consumers. The ability of PPA strains to infect eukaryotic cells should be tested to determine the risk level associated with edible plants.

### Plant-associated *P. aeruginosa* evolutionary relatedness to clinical isolates

16s rDNA-based phylogenetic tree revealed the evolutionary relatedness of the plant- and human-associated *P. aeruginosa*. Eighteen agricultural *P. aeruginosa* strains associated with cucumber, tomato, chili, and eggplant isolated in the current study, and ten strains from rice, guava, grass, pine, banana, lily, onion, ginseng, and aloe vera that were identified in previous studies clustered together with the human-pathogenic strains (Fig.3). Nearly 50% of these agricultural strains had more than 99% sequence similarity with the clinical isolate, PAO1. (Fig. 3). These results confirmed that plant- and human-associated *P. aeruginosa* are evolutionarily related. The current study supports the findings based on biochemical evidence by Schroth and his associates four decades ago (Schroth et al. 1977).

Phylogenetic analyses also showed that the agricultural *P. aeruginosa* strains do not always cluster by their plant source. The plant-associated *P. aeruginosa* strains retrieved from NCBI had high sequence similarity to the PPA strains isolated in this study (Fig. 3). An eggplant isolate, PPA12, co-clustered with a different strain SEGB6 (Accession no: MN565979) that was previously isolated from guava leaf. The two PPA strains, PPA08, and PPA15 had less than 97% DNA similarity with other *P. aeruginosa* strains, but their biochemical characteristics did not set them apart (Fig. 5-8). Whole-genome sequence analysis of these strains may reveal their novel characteristics. Nevertheless, the current study is the first attempt at exploring the evolutionary relationship of plant-associated *P. aeruginosa* with the well-characterized clinical isolates.

### *P. aeruginosa* strains did not cluster based on the plant source or niche

DNA fingerprint profiles revealed the molecular heterogeneity within the plant-associated *P. aeruginosa* strains (Fig. 4C). All the strains except for tomato isolates did not cluster based on the plant source or niche (rhizospheric and endophytic). For instance, each of the four strains, PPA15-PPA18 isolated from the chili plant, was associated with different clusters exposing their genetic variations (Fig. 4C). The occurrence of genomic variability among strains within a habitat was previously reported for clinical and environmental *P. aeruginosa* using ERIC fingerprinting (Martins et al. 2014).

In the current investigation, the phenotypic behaviors of the strains were also not associated with their plant source or niche. In the biochemical studies, the PPA strains from similar niches had varying abilities (Fig. 5-8). All the tomato-associated strains had similar DNA fingerprints but varied in phenotypic properties (Fig. 4C and 5D). In contrast, the chili-associated strains that had diverse fingerprints showed similar phenotypic traits and clustered together (Fig. 4C and 5D). One of the reasons for the lack of strong association by niche is that these crops are regularly replanted or rotated to preserve the soil health (Venter et al. 2016).

*P. aeruginosa* generally exhibits high levels of genomic plasticity and nearly 10-20% of its chromosome is comprised of accessory genes (Kung et al. 2010; Klockgether et al. 2011; Ozer et al. 2014; Pohl et al. 2014). This bacterium constantly evolves by acquiring new genetic segments from the environment through horizontal gene transfer (Shen et al. 2006; Mathee et al. 2008; Qiu et al. 2009; San Millan et al. 2015; Freschi et al. 2019). Multiple reports confirmed the within-patient patho-adaptive diversity of clinical *P. aeruginosa* (Wong et al. 2012; Jorth et al. 2015; Winstanley et al. 2016). The current study is the first of its kind to delineate the within-host genetic diversity of plant-associated *P. aeruginosa* isolates. Whole-genome sequencing and analysis would help to identify more precise genomic variations within these plant-associated strains.

## Concluding remarks

In the current study, we have shown the occurrence of *P. aeruginosa* strains in association with the vegetable plants including tomato, and cucumber that are often consumed raw. These plant-associated strains share comparable genetic and metabolic characteristics with the clinical isolates. It is likely that these strains are pathogenic, and it would be a serious health menace to the farmworkers and consumers if they breach the innate immune system. Future work will focus on further characterizing the isolates for their ability to cause disease using animal models. In addition, whole-genome sequence and analysis may reveal their total virulence potential.

## Acknowledgement

This work was not supported by any grant. SA was partially funded by the Fulbright Doctoral Nehru Research Fellowship by the U.S Department of State’s Bureau of Educational and Cultural Affairs and United-States India Educational Foundation (ID. PS00299273). SA also received Science and Engineering Research Board-International Travel Support funded by the Department of Science and Technology, India (No: ITS_2019_002449). We thank Dr. Sriyutha Murthy (Indira Gandhi Centre for Atomic Research, Kalpakkam, Tamilnadu, India) for providing *P. aeruginosa* strain PAO1.

## Author Contributions

The experiments were conceived and designed by SA and DB. The samples were processed by SA and PM, and the experiments were performed by SA. Critical analyses of the data were done by SA, KM and DB. The manuscript was prepared by SA and KM. Finally, all the authors were involved in the critical review of the paper.

## Ethical Approval

There was no human or animal subjects involved in this study.

## Data Availability

All sequence data generated in this study were deposited in NCBI GenBank (Accession no. MT734694 to MT734711).

## References

Abou Raji El Feghali, P. and Nawas, T. (2018) Pyocyanin: A powerful inhibitor of bacterial growth and biofilm formation. Madridge J Case Rep Stud 3, 101–107.

Adesemoye, A.O., Obini, M. and Ugoji, E.O. (2008) Comparison of plant growth-promotion with *Pseudomonas aeruginosa* and *Bacillus subtilis* in three vegetables. Braz J Microbiol 39, 423–426.

Adesemoye, A.O. and Ugoji, E.O. (2009) Evaluating *Pseudomonas aeruginosa* as plant growth-promoting rhizobacteria in West Africa. Arch Phytopathol Pflanzenschutz 42, 188–200.

Ahemad, M. and Khan, M.S. (2010) Phosphate-solubilizing and plant-growth-promoting *Pseudomonas aeruginosa* PS1 improves greengram performance in quizalafop-p-ethyl and clodinafop amended soil. Arch Environ Contam Toxicol 58, 361–372.

Akinsanya, M.A., Goh, J.K., Lim, S.P. and Ting, A.S.Y. (2015) Diversity, antimicrobial and antioxidant activities of culturable bacterial endophyte communities in *Aloe vera*. FEMS Microbiol Lett 362, fnv184.

Al-Kharousi, Z.S., Guizani, N., Al-Sadi, A.M., Al-Bulushi, I.M. and Shaharoona, B. (2016) Hiding in fresh fruits and vegetables: Opportunistic pathogens may cross geographical barriers. Int J Microbiol 2016, 4292417.

Alatraktchi, F.A.a., Svendsen, W.E. and Molin, S. (2020) Electrochemical detection of pyocyanin as a biomarker for *Pseudomonas aeruginosa*: A focused review. Sensors 20, 5218.

Ali, S., Charles, T.C. and Glick, B.R. (2017) Endophytic phytohormones and their role in plant growth promotion. In Functional Importance of the Plant Microbiome: Implications for Agriculture, Forestry and Bioenergy ed. Doty, S.L. pp.89–105. Cham: Springer International Publishing.

Ali Siddiqui, I. and Ehteshamul-Haque, S. (2001) Suppression of the root rot–root knot disease complex by *Pseudomonas aeruginosa* in tomato: The influence of inoculum density, nematode populations, moisture and other plant-associated bacteria. Plant Soil 237, 81–89.

Allydice-Francis, K. and Brown, P.D. (2012) Diversity of antimicrobial resistance and virulence determinants in *Pseudomonas aeruginosa* associated with fresh vegetables. Int J Microbiol 2012, 426241.

Alonso, A., Rojo, F. and Martínez, J.L. (1999) Environmental and clinical isolates of *Pseudomonas aeruginosa* show pathogenic and biodegradative properties irrespective of their origin. Environ Microbiol 1, 421–430.

Ambreetha, S. and Balachandar, D. (2019) Rhizobacteria-mediated root architectural improvement: A hidden potential for agricultural sustainability. In Plant Growth Promoting Rhizobacteria for Agricultural Sustainability : From Theory to Practices eds. Kumar, A. and Meena, V.S. pp.111–128. Singapore: Springer

Ambreetha, S., Chinnadurai, C., Marimuthu, P. and Balachandar, D. (2018) Plant-associated *Bacillus* modulates the expression of auxin-responsive genes of rice and modifies the root architecture. Rhizosphere 5, 57–66.

Anjaiah, V., Cornelis, P. and Koedam, N. (2003) Effect of genotype and root colonization in biological control of fusarium wilts in pigeonpea and chickpea by *Pseudomonas aeruginosa* PNA1. Can J Microbiol 49, 85–91.

Arif, M.S., Riaz, M., Shahzad, S.M., Yasmeen, T., Akhtar, M.J., Riaz, M.A., Jassey, V.E.J., Bragazza, L. and Buttler, A. (2016) Associative interplay of plant growth promoting rhizobacteria (*Pseudomonas aeruginosa* QS40) with nitrogen fertilizers improves sunflower (*Helianthus annuus* L.) productivity and fertility of aridisol. Appl Soil Ecol 108, 238–247.

Audenaert, K., Pattery, T., Cornelis, P. and Höfte, M. (2002) Induction of systemic resistance to *Botrytis cinerea* in tomato by P*seudomonas aeruginosa* 7NSK2: Role of salicylic acid, pyochelin, and pyocyanin. MPMI 15, 1147–1156.

Aznar, A., Chen, N.W.G., Rigault, M., Riache, N., Joseph, D., Desmaële, D., Mouille, G., Boutet, S., Soubigou-Taconnat, L., Renou, J.-P., Thomine, S., Expert, D. and Dellagi, A. (2014) Scavenging iron: A novel mechanism of plant immunity activation by microbial siderophores. Plant Physiol 164, 2167–2183.

Balamurugan, S. and Gunasekaran, S. (1996) Effect of combined inoculation of *Rhizoblum* sp and phosphobacteria at different levels of phosphorus in groundnut. Madras Agric J 83, 503–505.

Balasubramanian, D., Schneper, L., Kumari, H. and Mathee, K. (2012) A dynamic and intricate regulatory network determines *Pseudomonas aeruginosa* virulence. Nucleic Acids Res 41, 1–20.

Banerjee, A. and Dangar, T.K. (1995) *Pseudomonas aeruginosa*, a facultative pathogen of red palm weevil, *Rhynchophorus ferrugineus*. World J Microbiol Biotechnol 11, 618–620.

Bano, N. and Musarrat, J. (2003) Characterization of a new *Pseudomonas aeruginosa* strain nj-15 as a potential biocontrol agent. Curr Microbiol 46, 0324–0328.

Berger, C.N., Sodha, S.V., Shaw, R.K., Griffin, P.M., Pink, D., Hand, P. and Frankel, G. (2010) Fresh fruit and vegetables as vehicles for the transmission of human pathogens. Environ Microbiol 12, 2385–2397.

Botzenhart, K. and Doring, G. (1993) Ecology and epidemiology of *Pseudomonas aeruginosa*. In Pseudomonas aeruginosa as an opportunistic pathogen eds. Campa, M., Bendinelli, M. and Friedman, H. pp.1–18. Boston, MA: Springer US.

Bowya, T. and Balachandar, D. (2020) Harnessing PGPR inoculation through exogenous foliar application of salicylic acid and microbial extracts for improving rice growth. J Basic Microbiol 60, 950–961.

Bradbury, J.F. (1986) Guide to plant pathogenic bacteria. Kew, Surrey, UK: CAB International Mycological Institute.

Brindavathy, R. and Gopalaswamy, G. (2017) Effects of potassium releasing bacterium (KRB) with different levels of potash fertilizer in rice. Trend Biosci 10, 9193–9198.

Bunt, J.S. and Rovira, A.D. (1955) Microbiological studies of some subantarctic soils. J Soil Sci 6, 119–128.

Cappuccino, J.G. and Sherman, N. (1983) A laboratory manual. Addision*-*1999.

Cartwright, D.K., Chilton, W.S. and Benson, D.M. (1995) Pyrrolnitrin and phenazine production by *Pseudomonas cepacia*, strain 5.5B, a biocontrol agent of *Rhizoctonia solani*. Appl Microbiol Biotechnol 43, 211–216.

Chandra, H., Kumari, P., Bisht, R., Prasad, R. and Yadav, S. (2020) Plant growth promoting *Pseudomonas aeruginosa* from *Valeriana wallichii* displays antagonistic potential against three phytopathogenic fungi. Mol Biol Rep 47, 6015–6026.

Cho, J., JJ, C. and SD, K. (1975) Ornamental plants as carriers of *Pseudomonas aeruginosa*. Phytopathol 65, 425–431.

Clara, F. (1930) A new bacterial leaf disease of tobacco in the Philippines. Phytopathol 20, 691–706.

Clarke, K.R. (1993) Non-parametric multivariate analyses of changes in community structure. Aust J Ecol 18, 117–143.

Colombo, C., Palumbo, G., He, J.-Z., Pinton, R. and Cesco, S. (2014) Review on iron availability in soil: interaction of Fe minerals, plants, and microbes. J Soils Sediments 14, 538–548.

Correa, C.M.C., Tibana, A. and Filho, P.P.G. (1991) Vegetables as a source of infection with Pseudomonas aeruginosa in a University and Oncology Hospital of Rio de Janeiro. J Hosp Infect 18, 301–306.

Cother, E., Darbyshire, B. and Brewer, J. (1976) *Pseudomonas aerugisona*: Cause of internal brown rot of onion. Phytopathol 66, 828–834.

Curran, B., Morgan, J.A.W., Honeybourne, D. and Dowson, C.G. (2005) Commercial mushrooms and bean sprouts are a source of *Pseudomonas aeruginosa*. J Clin Microbiol 43, 5830–5831.

De Meyer, G. and Höfte, M. (1997) Salicylic acid produced by the rhizobacterium *Pseudomonas aeruginosa* 7nsk2 induces resistance to leaf infection by *Botrytis cinerea* on bean. Phytopathol 87, 588–593.

De Vleesschauwer, D., Cornelis, P. and Höfte, M. (2006) Redox-active pyocyanin secreted by *Pseudomonas aeruginosa* 7NSK2 triggers systemic resistance to *Magnaporthe grisea* but enhances *Rhizoctonia solani* Susceptibility in rice. MPMI 19, 1406–1419.

Deredjian, A., Colinon, C., Hien, E., Brothier, E., Youenou, B., Cournoyer, B., Dequiedt, S., Hartmann, A., Jolivet, C., Houot, S., Ranjard, L., Saby, N.P. and Nazaret, S. (2014) Low occurrence of *Pseudomonas aeruginosa* in agricultural soils with and without organic amendment. Front Cell Infect Microbiol 4, 53.

Devi, K.A., Pandey, G., Rawat, A.K.S., Sharma, G.D. and Pandey, P. (2017) The endophytic symbiont—*Pseudomonas aeruginosa* stimulates the antioxidant activity and growth of *Achyranthes aspera L*. Front Microbiol 8, 1–14.

Devnath, P., Uddin, M.K. and Aha, F. (2017) Extraction, purification and charact *Pseudomonas aeruginosa*. Int Res J Biol Sci 6, 1–9.

Doustdar, F., Karimi, F., Abedinyfar, Z., Amoli, F.A. and Goudarzi, H. (2019) Genetic features of *Pseudomonas aeruginosa* isolates associated with eye infections referred to Farabi Hospital, Tehran, Iran. Int Ophthalmol 39, 1581–1587.

Durairaj, K., Velmurugan, P., Park, J.-H., Chang, W.-S., Park, Y.-J., Senthilkumar, P., Choi, K.-M., Lee, J.-H. and Oh, B.-T. (2017) Potential for plant biocontrol activity of isolated *Pseudomonas aeruginosa* and *Bacillus stratosphericus* strains against bacterial pathogens acting through both induced plant resistance and direct antagonism. FEMS Microbiol Lett 364, fnx225.

Edi-Premono, M., Moawad and PLG, V. (1996) Effect of phosphate solubilizing *Pseudmonas putida* on the growth of maize and its survival in the rhizosphere. Indones J Crop Sci 11, 13–23.

El-Said, H., Gewaily, E., Salem, S. and Tohami, M. (1982) Biological and chemical control of *Pseudomonas aeruginosa*, the causal organism of blight disease of bean plant in Egypt. Egypt J Microbiol 17, 65–80.

Elbadry, M., El-Bassel, A. and Elbanna, K. (1999) Occurrence and dynamics of phototrophic purple nonsulphur bacteria compared with other asymbiotic nitrogen fixers in ricefields of Egypt. World J Microbiol Biotechnol 15, 359–362.

Elrod, R.P. and Braun, A.C. (1942) *Pseudomonas aeruginosa*: Its rôle as a plant pathogen. J Bacteriol 44, 633–645.

Essar, D.W., Eberly, L., Hadero, A. and Crawford, I.P. (1990) Identification and characterization of genes for a second anthranilate synthase in *Pseudomonas aeruginosa*: interchangeability of the two anthranilate synthases and evolutionary implications. J Bacteriol 172, 884–900.

Fasim, F., Ahmed, N., Parsons, R. and Gadd, G.M. (2002) Solubilization of zinc salts by a bacterium isolated from the air environment of a tannery. FEMS Microbiol Lett 213, 1–6.

Freschi, L., Vincent, A.T., Jeukens, J., Emond-Rheault, J.-G., Kukavica-Ibrulj, I., Dupont, M.-J., Charette, S.J., Boyle, B. and Levesque, R.C. (2019) The *Pseudomonas aeruginosa* pan-genome provides new insights on its population structure, horizontal gene transfer, and pathogenicity. Genome Biol Evol 11, 109–120.

Gao, J., Wang, Y., Wang, C.W. and Lu, B.H. (2014) First report of bacterial root rot of ginseng caused by *Pseudomonas aeruginosa* in China. Plant Dis 98, 1577–1577.

Gardner, J.M., Feldman, A.W. and Zablotowicz, R.M. (1982) Identity and behavior of xylem-residing bacteria in rough lemon roots of florida citrus trees. Appl Environ Microbiol 43, 1335–1342.

Gessard, C. (1984) On the blue and green coloration that appears on bandages. Rev Infect Dis 6, S775–S776.

Ghosh, S.K., Bera, T. and Chakrabarty, A.M. (2020) Microbial siderophore – A boon to agricultural sciences. Biol Control 144, 104214.

Gomathy, M., Thangaraju, M., Gunasekaran, S. and Gopal, N.O. (2007) Sporulation and regeneration efficiency of phosphobacteria *(Bacillus megaterium* var *phosphaticum*). Indian J Microbiol 47, 259–262.

Gordon, S.A. and Weber, R.P. (1951) Colorimetric estimation of indoleacetic acid. Plant Physiol 26, 192–195.

Goswami, D., Dhandhukia, P., Patel, P. and Thakker, J.N. (2014) Screening of PGPR from saline desert of Kutch: Growth promotion in *Arachis hypogea* by *Bacillus licheniformis* A2. Microbiol Res 169, 66–75.

Green, S.K., Schroth, M.N., Cho, J.J., Kominos, S.D. and Vitanza-Jack, V.B. (1974) Agricultural plants and soil as a reservoir for *Pseudomonas aeruginosa*. Appl Microbiol 28, 987–991.

Gupta, R., Usha, M. and Pandey, U. (1986) Bacterial brown rot of onion in storage-a new record from India. Indian J Plant Pathol 4, 184.

Gupta, V. and Buch, A. (2019) *Pseudomonas aeruginosa* predominates as multifaceted rhizospheric bacteria with combined abilities of P-solubilization and biocontrol. J Pure Appl Microbiol 13, 319–328.

Hall, S., McDermott, C., Anoopkumar-Dukie, S., McFarland, A.J., Forbes, A., Perkins, A.V., Davey, A.K., Chess-Williams, R., Kiefel, M.J., Arora, D. and Grant, G.D. (2016) Cellular effects of pyocyanin, a secreted virulence factor of *Pseudomonas aeruginosa*. Toxins 8, 236–244.

Hariprasad, P., Chandrashekar, S., Singh, S.B. and Niranjana, S.R. (2014) Mechanisms of plant growth promotion and disease suppression by *Pseudomonas aeruginosa* strain 2apa. J Basic Microbiol 54, 792–801.

Haynes, W.C. (1951) *Pseudomonas aeruginosa*---its characterization and identification. Microbiol 5, 939–950.

He, J., Baldini, R.L., Déziel, E., Saucier, M., Zhang, Q., Liberati, N.T., Lee, D., Urbach, J., Goodman, H.M. and Rahme, L.G. (2004) The broad host range pathogen *Pseudomonas aeruginosa* strain PA14 carries two pathogenicity islands harboring plant and animal virulence genes. PNAS 101, 2530–2535.

Holloway, B.W. (1955) Genetic recombination in *Pseudomonas aeruginosa*. J Gen Microbiol 13, 572–581.

Hu, X., Chen, J. and Guo, J. (2006) Two phosphate- and potassium-solubilizing bacteria isolated from Tianmu Mountain, Zhejiang, China. World J Microbiol Biotechnol 22, 983–990.

Iasur Kruh, L., Bari, V.K., Abu-Nassar, J., Lidor, O. and Aly, R. (2020) Characterization of an endophytic bacterium (*Pseudomonas aeruginosa*), originating from tomato (*Solanum lycopersicum* L.), and its ability to inhabit the parasitic weed *Phelipanche aegyptiaca*. Plant Signal Behav 15, 1766292.

Illmer, P. and Schinner, F. (1992) Solubilization of inorganic phosphates by microorganisms isolated from forest soils. Soil Biol Biochem 24, 389–395.

Ingledew, W.M. and Campbell, J.J. (1969) A new resuspension medium for pyocyanine production. Can J Microbiol 15, 595–598.

Janssen, P.H. (2006) Identifying the dominant soil bacterial taxa in libraries of 16S rRNA and 16S rRNA Genes. Appl Environ Microbiol 72, 1719–1728.

Jasim, B., Anisha, C., Rohini, S., Kurian, J.M., Jyothis, M. and Radhakrishnan, E.K. (2014) Phenazine carboxylic acid production and rhizome protective effect of endophytic *Pseudomonas aeruginosa* isolated from *Zingiber officinale*. World J Microbiol Biotechnol 30, 1649–1654.

Jha, B.K., Gandhi Pragash, M., Cletus, J., Raman, G. and Sakthivel, N. (2009) Simultaneous phosphate solubilization potential and antifungal activity of new fluorescent pseudomonad strains, *Pseudomonas aeruginosa*, *P. plecoglossicida* and *P. mosselii*. World J Microbiol Biotechnol 25, 573–581.

Jorth, P., Staudinger, Benjamin J., Wu, X., Hisert, K.B., Hayden, H., Garudathri, J., Harding, Christopher L., Radey, Matthew C., Rezayat, A., Bautista, G., Berrington, William R., Goddard, Amanda F., Zheng, C., Angermeyer, A., Brittnacher, Mitchell J., Kitzman, J., Shendure, J., Fligner, Corinne L., Mittler, J., Aitken, Moira L., Manoil, C., Bruce, James E., Yahr, Timothy L. and Singh, Pradeep K. (2015) Regional isolation drives bacterial diversification within cystic fibrosis lungs. Cell Host Microbe 18, 307–319.

Kim, B.S., Lee, J.Y. and Hwang, B.K. (2000) *In vivo* control and in vitro antifungal activity of rhamnolipid B, a glycolipid antibiotic, against *Phytophthora capsici* and *Colletotrichum orbiculare*. Pest Manag Sci 56, 1029–1035.

King, E.O., Ward, M.K. and Raney, D.E. (1954) Two simple media for the demonstration of pyocyanin and fluorescin. J Lab Clin Med 44, 301–307.

Klockgether, J., Cramer, N., Wiehlmann, L., Davenport, C. and Tümmler, B. (2011) *Pseudomonas aeruginosa* genomic structure and diversity. Front Microbiol 2.

Kloepper, J.W., Leong, J., Teintze, M. and Schroth, M.N. (1980) *Pseudomonas siderophores:* A mechanism explaining disease-suppressive soils. Curr Microbiol 4, 317–320.

Kominos, S.D., Copeland, C.E., Grosiak, B. and Postic, B. (1972) Introduction of *Pseudomonas aeruginosa* into a hospital via vegetables. Appl Microbiol 24, 567–570.

Kravchenko, L.V., Azarova, T.S., Makarova, N.M. and Tikhonovich, I.A. (2004) The effect of tryptophan present in plant root exudates on the phytostimulating activity of Rhizobacteria. Microbiol 73, 156–158.

Kumar, A., Munder, A., Aravind, R., Eapen, S.J., Tümmler, B. and Raaijmakers, J.M. (2013) Friend or foe: genetic and functional characterization of plant endophytic *Pseudomonas aeruginosa*. Environ Microbiol 15, 764–779.

Kumar, A.S., Sridar, R. and Uthandi, S. (2017) Mitigation of drought in rice by a phyllosphere bacterium *Bacillus altitudinis* FD48. Afr J Microbiol Res 11, 1614–1625.

Kung, V.L., Ozer, E.A. and Hauser, A.R. (2010) The accessory genome of *Pseudomonas aeruginosa*. MMBR 74, 621–641.

Kurachi, M. (1958) Studies on the biosynthesis of pyocyanine.(II): Isolation and determination of pyocyanine. *Bulletin of the Institute for Chemical Research, Kyoto University* 36, 174–187.

Lebeda, A., Kudela, V. and Jedlickova, Z. (1984) Pathogenicity of *Pseudomonas aeruginosa* for plants and animals. Acta Phytopathol Acad Sci Hung 19, 271–284.

Leong, J. (1986) Siderophores: their biochemistry and possible role in the biocontrol of plant pathogens. Annu Rev Phytopathol 24, 187–209.

Linu, M.S., Asok, A.K., Thampi, M., Sreekumar, J. and Jisha, M.S. (2019) Plant growth promoting traits of indigenous phosphate solubilizing *Pseudomonas aeruginosa* isolates from chilli (*Capsicumannuum* L.) rhizosphere. Commun Soil Sci Plant Anal 50, 444–457.

Mahmoud, S.Y., Ziedan, E.-S.H., Farrag, E.S., Kalafalla, R.S. and Ali, A.M. (2016) Antifungal activity of pyocyanin produced by *Pseudomonas aeruginosa* against *Fusarium oxysporum* Schlech phytopathogenic fungi. Int J PharmTech Res 9, 43–50.

Marathe, R., Phatake, Y., Shaikh, A., Shinde, B. and Gajbhiye, M. (2017) Effect of IAA produced by *Pseudomonas aeruginosa* 6a (bc4) on seed germination and plant growth of *Glycin max*. J Exp Biol Agric Sci 5, 351–358.

Martins, V.V., Pitondo-Silva, A., de Melo Manço, L., Falcão, J.P., Freitas, S.d.S., da Silveira, W.D. and Stehling, E.G. (2014) Pathogenic potential and genetic diversity of environmental and clinical isolates of *Pseudomonas aeruginosa*. APMIS 122, 92–100.

Mathee, K. (2018) Forensic investigation into the origin of *Pseudomonas aeruginosa* PA14 — old but not lost. J Med Microbiol 67, 1019–1021.

Mathee, K., Narasimhan, G., Valdes, C., Qiu, X., Matewish, J.M., Koehrsen, M., Rokas, A., Yandava, C.N., Engels, R., Zeng, E., Olavarietta, R., Doud, M., Smith, R.S., Montgomery, P., White, J.R., Godfrey, P.A., Kodira, C., Birren, B., Galagan, J.E. and Lory, S. (2008) Dynamics of *Pseudomonas aeruginosa* genome evolution. PNAS 105, 3100–3105.

Melody, S.C. (1997) Plant Molecular Biology - A laboratory manual. New York, USA: Springer-Verlag Berlin Heidelberg.

Mondal, K.K., Mani, C., Singh, J., Dave, S.R., Tipre, D.R., Kumar, A. and Trivedi, B.M. (2012) Fruit rot of tinda caused by *Pseudomonas aeruginosa*–A new report from India. Plant Dis 96, 141–141.

Moradali, M.F., Ghods, S. and Rehm, B.H.A. (2017) *Pseudomonas aeruginosa* lifestyle: A paradigm for adaptation, survival, and persistence. Front Cell Infect Microbiol 7, 1–29.

Mukherjee, A., Singh, B.K. and Verma, J.P. (2020) Harnessing chickpea (*Cicer arietinum* L.) seed endophytes for enhancing plant growth attributes and bio-controlling against *Fusarium* sp. Microbiol Res 237, 126469.

Narayanasamy, S., Thangappan, S. and Uthandi, S. (2020) Plant growth-promoting *Bacillus* sp. cahoots moisture stress alleviation in rice genotypes by triggering antioxidant defense system. Microbiol Res 239, 126518.

Nithya, A. and Babu, S. (2017) Prevalence of plant beneficial and human pathogenic bacteria isolated from salad vegetables in India. BMC Microbiol 17, 64.

Noura, Salih, K.M., Jusuf, N.H., Hamid, A.A. and Yusoff, W.M.W. (2009) High prevalence of *Pseudomonas* species in soil samples from Ternate Island-Indonesia. Pak J Biol Sci 12, 1036–1040.

Obaton, M., Amarger, N. and Alexander, M. (1968) Heterotrophic nitrification by *Pseudomonas aeruginosa*. Arch Microbiol 63, 122–132.

Oleńska, E., Małek, W., Wójcik, M., Swiecicka, I., Thijs, S. and Vangronsveld, J. (2020) Beneficial features of plant growth-promoting rhizobacteria for improving plant growth and health in challenging conditions: A methodical review. Sci Total Environ 743, 140682.

Ozer, E.A., Allen, J.P. and Hauser, A.R. (2014) Characterization of the core and accessory genomes of *Pseudomonas aeruginosa* using bioinformatic tools Spine and AGEnt. BMC Genomics 15, 737.

Picard, B., Denamur, E., Barakat, A., Elion, J. and Goullet, P. (1994) Genetic heterogeneity of *Pseudomonas aeruginosa* clinical isolates revealed by esterase electrophoretic polymorphism and restriction fragment length polymorphism of the ribosomal RNA gene region. J Med Microbiol 40, 313–322.

Pohl, S., Klockgether, J., Eckweiler, D., Khaledi, A., Schniederjans, M., Chouvarine, P., Tümmler, B. and Häussler, S. (2014) The extensive set of accessory *Pseudomonas aeruginosa* genomic components. FEMS Microbiol Lett 356, 235–241.

Priyaja, P., Jayesh, P., Philip, R. and Singh, I.B. (2016) Pyocyanin induced in vitro oxidative damage and its toxicity level in human, fish and insect cell lines for its selective biological applications. Cytotechnology 68, 143–155.

Qiu, X., Kulasekara, B. and Lory, S. (2009) Role of horizontal gene transfer in the evolution of Pseudomonas aeruginosa virulence. In Microbial Pathogenomics. pp.126–139: Karger Publishers.

Radford, R., Brahma, A., Armstrong, M. and Tullo, A.B. (2000) Severe sclerokeratitis due to *Pseudomonas aeruginosa* in non-contact-lens wearers. Eye 14, 3–7.

Radhapriya, P., Ramachandran, A., Anandham, R. and Mahalingam, S. (2015) *Pseudomonas aeruginosa* RRALC3 enhances the biomass, nutrient and carbon contents of pongamia pinnata seedlings in degraded forest soil. Plos One 10, e0139881.

Rahme, L., Stevens, E., Wolfort, S., Shao, J., Tompkins, R. and Ausubel, F. (1995) Common virulence factors for bacterial pathogenicity in plants and animals. Science 268, 1899–1902.

Rahme, L.G., Ausubel, F.M., Cao, H., Drenkard, E., Goumnerov, B.C., Lau, G.W., Mahajan-Miklos, S., Plotnikova, J., Tan, M.-W., Tsongalis, J., Walendziewicz, C.L. and Tompkins, R.G. (2000) Plants and animals share functionally common bacterial virulence factors. PNAS 97, 8815–8821.

Reyes, E.A., Bale, M.J., Cannon, W.H. and Matsen, J.M. (1981) Identification of *Pseudomonas aeruginosa* by pyocyanin production on Tech agar. J Clin Microbiol 13, 456–458.

Reynolds, H.Y., Levine, A.S., Wood, A.E., Zierdt, C.H., Dale, D.C. and E., P. (1975) *Pseudomonas aeruginosa* infections: Persisting problems and current research to find new therapies. Ann Intern Med 82, 819–831.

Ribeiro, I.D.A., Volpiano, C.G., Vargas, L.K., Granada, C.E., Lisboa, B.B. and Passaglia, L.M.P. (2020) Use of mineral weathering bacteria to enhance nutrient availability in crops: A review. Front Plant Sci 11, 590774–590774.

Rosenthal, V.D., Bat-Erdene, I., Gupta, D., Belkebir, S., Rajhans, et al. (2020) International Nosocomial Infection Control Consortium (INICC) report, data summary of 45 countries for 2012-2017: Device-associated module. Am J Infect 48, 423–432.

Roychowdhury, R., Qaiser, T.F., Mukherjee, P. and Roy, M. (2019) Isolation and characterization of a *Pseudomonas aeruginosa* strain PGP for plant growth promotion. *Proc Natl Acad Sci India*, Sect B Biol Sci 89, 353–360.

Sah, S., Singh, N. and Singh, R. (2017) Iron acquisition in maize (*Zea mays* L.) using *Pseudomonas* siderophore. 3 Biotech 7, 121–121.

Saitou, N. and Nei, M. (1987) The neighbor-joining method: a new method for reconstructing phylogenetic trees. Mol Biol Evol 4, 406–425.

San Millan, A., Toll-Riera, M., Qi, Q. and MacLean, R.C. (2015) Interactions between horizontally acquired genes create a fitness cost in *Pseudomonas aeruginosa*. Nat Commun 6, 6845.

Sanger, F., Nicklen, S. and Coulson, A.R. (1977) DNA sequencing with chain-terminating inhibitors. PNAS 74, 5463–5467.

Schroth, M., Cho, J., Green, S. and Kominos, S. (1977) Epidemiology of *Pseudomonas aeruginosa* in agricultural areas [Soil and plants serve as the natural and permanent reservoirs for the bacterium]. In Pseudomonas aeruginosa: Ecological aspects and patient colonization ed. Young, V. pp.1–29: New York: Raven Press.

Schroth, M.N., Cho, J.J., Green, S.K., Kominos, S.D. and Publishing, M.S. (2018) Epidemiology of *Pseudomonas aeruginosa* in agricultural areas. J Med Microbiol 67, 1191–1201.

Schwyn, B. and Neilands, J.B. (1987) Universal chemical assay for the detection and determination of siderophores. Anal Biochem 160, 47–56.

Shanmugaiah, V., Mathivanan, N. and Varghese, B. (2010) Purification, crystal structure and antimicrobial activity of phenazine-1-carboxamide produced by a growth-promoting biocontrol bacterium, *Pseudomonas aeruginosa* MML2212. J Appl Microbiol 108, 703–711.

Shen, K., Sayeed, S., Antalis, P., Gladitz, J., Ahmed, A., Dice, B., Janto, B., Dopico, R., Keefe, R., Hayes, J., Johnson, S., Yu, S., Ehrlich, N., Jocz, J., Kropp, L., Wong, R., Wadowsky, R.M., Slifkin, M., Preston, R.A., Erdos, G., Post, J.C., Ehrlich, G.D. and Hu, F.Z. (2006) Extensive genomic plasticity in *Pseudomonas aeruginosa* revealed by identification and distribution studies of novel genes among clinical isolates. Infect Immun 74, 5272–5283.

Shi, Z., Ren, D., Hu, S., Hu, X., Wu, L., Lin, H., Hu, J., Zhang, G. and Guo, L. (2015) Whole genome sequence of *Pseudomonas aeruginosa* F9676, an antagonistic bacterium isolated from rice seed. J Biotechnol 211, 77–78.

Singh, P., Singh, R.K., Guo, D.J., Sharma, A., Singh, R.N., Li, D.P., Malviya, M.K., Song, X.P., Lakshmanan, P., Yang, L.T. and Li, Y.R. (2021) Whole genome analysis of sugarcane root-associated endophyte *Pseudomonas aeruginosa* B18-A plant growth-promoting bacterium with antagonistic potential against *Sporisorium scitamineum*. Front Microbiol 12, 628376.

Sperber, J. (1958) Solution of apatite by soil microorganisms producing organic acids. Aust J Agr Res 9, 782–787.

Spilker, T., Coenye, T., Vandamme, P. and LiPuma, J.J. (2004) PCR-based assay for differentiation of *Pseudomonas aeruginosa* from other pseudomonas species recovered from cystic fibrosis patients. J Clin Microbiol 42, 2074–2079.

Sukumar, p., legué, v., vayssières, a., martin, f., tuskan, g.a. and Kalluri, u.c. (2013) Involvement of auxin pathways in modulating root architecture during beneficial plant–microorganism interactions. Plant Cell Environ 36, 909–919.

Sulochana, M.B., Jayachandra, S.Y., Kumar, S.A., Parameshwar, A.B., Reddy, K.M. and Dayanand, A. (2014) Siderophore as a potential plant growth-promoting agent produced by *Pseudomonas aeruginosa* JAS-25. Appl Biochem Biotechnol 174, 297–308.

Sun, X., Xu, Y., Chen, L., Jin, X. and Ni, H. (2021) The salt-tolerant phenazine-1-carboxamide-producing bacterium *Pseudomonas aeruginosa* NF011 isolated from wheat rhizosphere soil in dry farmland with antagonism against *Fusarium graminearum*. Microbiol Res 245, 126673.

Tamura, K., Nei, M. and Kumar, S. (2004) Prospects for inferring very large phylogenies by using the neighbor-joining method. PNAS 101, 11030–11035.

Tate, D., Mawer, S. and Newton, A. (2003) Outbreak of *Pseudomonas aeruginosa* folliculitis associated with a swimming pool inflatable. Epidemiol Infect 130, 187–192.

Tiwari, P. and Singh, J.S. (2017) A plant growth promoting rhizospheric pseudomonas aeruginosa strain inhibits seed germination in *Triticum aestivum* (L) and *Zea mays* (L). Microbiol Res 8, 7233.

Turner, J.M. and Messenger, A.J. (1986) Occurrence, Biochemistry and Physiology of Phenazine Pigment Production. In Advances in Microbial Physiology eds. Rose, A.H. and Tempest, D.W. pp.211–275: Academic Press.

Van Soestbergen, A.A. and Lee, C.H. (1969) Pour plates or streak plates? Appl Microbiol 18, 1092–1093.

Venter, Z.S., Jacobs, K. and Hawkins, H.-J. (2016) The impact of crop rotation on soil microbial diversity: A meta-analysis. Pedobiologia 59, 215–223.

Versalovic, J., Schneider, M., De Bruijn, F.J. and Lupski, J.R. (1994) Genomic fingerprinting of bacteria using repetitive sequence-based polymerase chain reaction. Method Mol Cell Biol 5, 25–40.

Viswanathan, P. and Kaur, R. (2001) Prevalence and growth of pathogens on salad vegetables, fruits and sprouts. Int J Hyg Environ Health 203, 205–213.

Von Graevenitz, A. (1977) The role of opportunistic bacteria in human disease. Annu Rev Microbiol 31, 447–471.

Waksman, S.A. and Woodruff, H.B. (1940) The soil as a source of microorganisms antagonistic to disease-producing bacteria. J Bacteriol 40, 581–600.

Weisburg, W.G., Barns, S.M., Pelletier, D.A. and Lane, D.J. (1991) 16S ribosomal DNA amplification for phylogenetic study. J Bacteriol 173, 697–703.

Wilson, L.A. and Sharp, P.M. (2006) Enterobacterial Repetitive Intergenic Consensus (ERIC) sequences in *Escherichia coli*: Evolution and implications for ERIC-PCR. Mol Biol Evol 23, 1156–1168.

Winsor, G.L., Griffiths, E.J., Lo, R., Dhillon, B.K., Shay, J.A. and Brinkman, F.S. (2016) Enhanced annotations and features for comparing thousands of Pseudomonas genomes in the Pseudomonas genome database. Nucleic Acids Res 44, D646–653.

Winstanley, C., O’Brien, S. and Brockhurst, M.A. (2016) *Pseudomonas aeruginosa* evolutionary adaptation and diversification in cystic fibrosis chronic lung infections. Trends Microbiol 24, 327–337.

Wong, A., Rodrigue, N. and Kassen, R. (2012) Genomics of Adaptation during experimental evolution of the opportunistic pathogen *Pseudomonas aeruginosa*. PLOS Genetics 8, e1002928.

Wright, C., Kominos, S.D. and Yee, R.B. (1976) Enterobacteriaceae and *Pseudomonas aeruginosa* recovered from vegetable salads. Appl Environ Microbiol 31, 453–454.

Wu, T., Xu, J., Xie, W., Yao, Z., Yang, H., Sun, C. and Li, X. (2018) *Pseudomonas aeruginosa* L10: A hydrocarbon-degrading, biosurfactant-producing, and plant-growth-promoting endophytic bacterium isolated from a reed (*Phragmites australis*). Front Microbiol 9, 1087.

Yasmin, S., Hafeez, F.Y. and Rasul, G. (2014) Evaluation of *Pseudomonas aeruginosa* Z5 for biocontrol of cotton seedling disease caused by *Fusarium oxysporum*. BioControl Sci Techn 24, 1227–1242.

Zhao, Y. (2010) Auxin biosynthesis and its role in plant development. Annu Rev Plant Biol 61, 49–64.

